# Oscillations in working memory and neural binding: a mechanism for multiple memories and their interactions

**DOI:** 10.1101/263277

**Authors:** Jason E. Pina, Mark Bodner, Bard Ermentrout

## Abstract

Neural oscillations have been implicated in many different basic brain and cognitive processes. Oscillatory activity has been suggested to play a role in neural binding, and more recently in the maintenance of information in working memory. This latter work has focused primarily on oscillations in terms of providing a “code” in working memory. However, oscillations may additionally play a fundamental role in essential properties and behaviors that neuronal networks must exhibit in order to produce functional working memory. In the present work, we present a biologically plausible working memory model and demonstrate that specific types of stable oscillatory dynamics may play a critical role in facilitating properties of working memory, including transitions between different memory states and a multi-item working memory capacity. We also show these oscillatory dynamics may facilitate and provide an underlying mechanism to enable a range of different types of binding in the context of working memory.

**Author summary:** Working memory is a form of short-term memory that is limited in capacity to perhaps 3 – 5 items. Various studies have shown that ensembles of neurons oscillate during working memory retention, and cross-frequency coupling (between, e.g., theta and gamma frequencies) has been conjectured as underlying the observed limited capacity. Binding occurs when different objects or concepts are associated with each other and can persist as working memory representations; neuronal synchrony has been hypothesized as the neural correlate. We propose a novel computational model of a network of oscillatory neuronal populations that capture salient attributes of working memory and binding by allowing for both stable synchronous and asynchronous activity. The oscillatory dynamics we describe may provide a mechanism that can facilitate aspects of working memory, such as maintaining multiple items active at once, creating rich neural representations of memories via binding, and rapidly transitioning activtation patterns based on selective inputs.

## 1 Introduction

The persistent elevated firing rates of populations of neurons concomitant with the maintenance of information associated with memoranda has been established as the neuronal substrate of working memory, and numerous physiological and imaging studies have verified these dynamics at the single neuron, population, and network levels [1–6]. To function properly, however, these neurons, populations and networks must be able to be activated rapidly by external or internal input corresponding to aspects of memoranda, that activation must be selective and stable, and the elevated firing rates must be able to return to background levels when the information is no longer needed. Furthermore, the requirements of cognition and behavior imply that mechanisms must exist for rapidly transitioning among sequences of active memories, and that multiple (and possibly overlapping) selective populations and networks can be simultaneously active. This applies both to the case of neural binding, in which neuronal populations or networks (e.g., corresponding to different aspects of a particular memorandum or working memory task) must be combined and maintained, and for the case in which multiple different memoranda are simultaneously maintained in working memory. In this latter case, there are well established approximate upper limits to the number of separate memoranda that can be simultaneously maintained. Indeed, one of the fundamental properties of working memory is that it has a limited capacity, possibly limited to three to five objects [7–11].

In its most general terms, binding refers to how items encoded in distinct brain circuits or neural populations can be combined for perception, decisions, and action, and has been partitioned to encompass multiple situations [12]. These different binding types include feature binding, which involves the association of different characteristics to an object and how we make and then unravel these associations, and variable binding, which arises, for example, in language and other symbolic thought (e.g., the binding of a variable name and its value). In every instance, some form of synchronization of neural activity has been proposed as the underlying mechanism (e.g., [12–18]). Binding is a key aspect in working memory, as most objects we encounter, physical or symbolic, are multi-featured. There is also a limited capacity to the number of features that may be represented for objects; however, it is unclear at present exactly how this feature capacity relates to the aforementioned working memory capacity [9, 19, 20].

A major type of dynamics that has been identified with cognitive function in general, and increasingly with working memory in particular, is oscillatory in nature (e.g., [21–23]). While the presence of different patterns of oscillations is well documented, the specific roles they play are not well understood. Recent work has suggested, however, that oscillations in various frequency bands and coupled or nested oscillations could play a fundamental role in the functioning of different aspects of working memory [6, 24–27]. Much of this work has focused on the role of oscillatory states in representing a “code” of working memory. However, oscillatory dynamics may also play a critical generic role in facilitating the range of dynamics and optimized conditions required for working memory function as described above.

In the present work, we investigate the role of oscillatory dynamics in working memory network models with plausible architectures and parameter values in carrying out or facilitating these critical functions. In particular, we examine the role oscillatory states can play as an underlying mechanism to allow for multiple stable, active memoranda, to establish binding via synchronous relationships, to transition between different active working memory states, and to rapidly activate and terminate activity in those networks as required by the needs of cognition and thought. We show how oscillatory dynamics may facilitate these potentially competing requirements, and identify and discuss critical network parameters involved in achieving these dynamics.

## 2 Model and methods

Oscillations during memory task delays have been seen in population-level activity, including in local field potential recordings and human EEG traces [4, 5]. This motivates us to model working memory oscillatory dynamics at the ensemble level, using Wilson-Cowan type equations for the firing rates (or synaptic activities) of each local circuit that will comprise the model. Since previous experimental and computational studies suggest NMDA receptors may be crucial to the persistent elevated firing rates associated with proper working memory function in experimental and computational studies, we model the effect of NMDA receptors as a separate component [2, 28–31]:

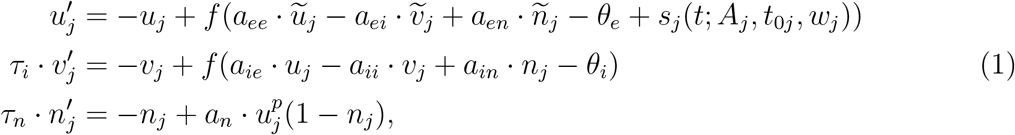

with *j* ∈ {1, …, *N*}, where *N* is the number of interconnected populations, or *u* − *v* − *n* triplets. Here, *u*_*j*_ represents the fast AMPA excitatory synapses in the population, *v*_*j*_ the slower inhibition, and *n*_*j*_ a slow NMDA component of the excitatory synapse. We assume coupled populations reside in neighboring areas, so that delays need not be considered, and that *τ*_*n*_ > *τ*_*i*_ > 1, where we have rescaled time so that the timescale is 1 for the fast excitation. Fig 1A shows a schematic of the connectivities. Since we do not explicitly model membrane potential or spiking, there is no voltage dependence to the NMDA component; rather, it acts as a slow excitatory input. The NMDA component also saturates at 1, while the faster variables, *u* and *v*, are not constrained. The connection strength from *x* to *y* is given by the coefficient *a*_*yx*_. The parameters *a*_*n*_ and *p* in the NMDA equation allow us to adjust the average NMDA level when that component is active; larger *p* also results in faster NMDA activation since *u* is typically larger than 1 when activated. Note that the behavior is fairly insensitive to *p*; the *n* − *u* null surface (where *n*’ = 0) is sigmoidal and *p* increases the gain of this sigmoid.

**Fig 1.**
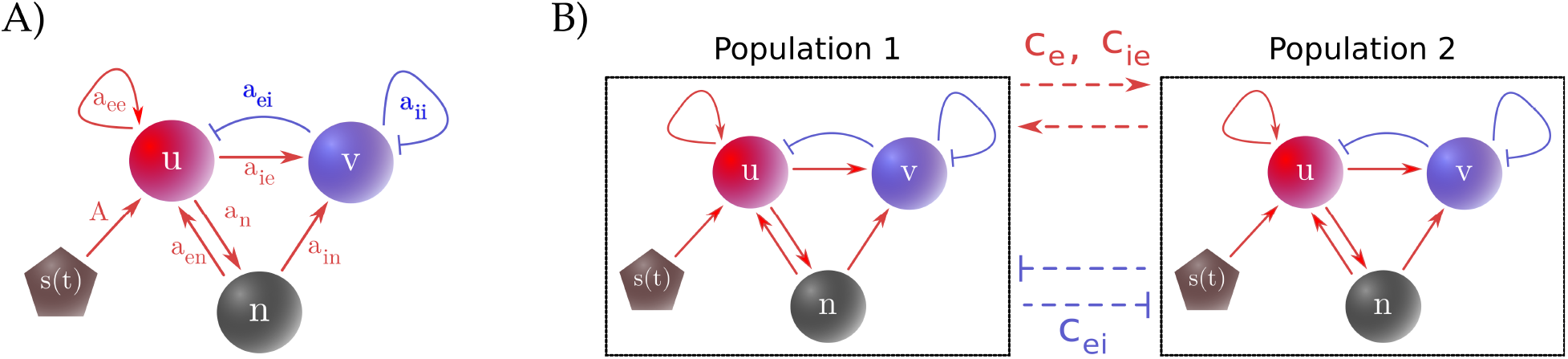
Model connectivity for one and two populations. (A) One population as described in Eq (1). *u* is the fast excitatory AMPA synaptic activity, *v* the inhibitory GABA activity, and *n* the slow excitatory NMDA activity. Feedforward excitation to the AMPA synapses (*u*) triggers activity in the system. (B) Connectivity for two populations, each with three components as in (A). The populations are connected with excitatory and inhibitory coupling as described in Eqs (2) and (3).

An input stimulus, *s*_*j*_ (*t*), allows the system to be activated. This stimulus is a square wave with onset time 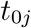, width *w*_*j*_, and amplitude *A*_*j*_. For one population (*N* = 1), 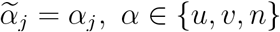; this will change as we add more populations. The firing rate is given by 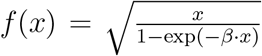. We have chosen this firing rate since it gives an approximation of the firing rate of a noisy quadratic-integrate-and-fire spiking neuron [32, Chapter 10]. Additionally, in Appendix B, we show an example simulation of a spiking model and compare it to the one-population version of the Wilson-Cowan equations.

Each *u* – *v* – *n* triplet represents a tightly recurrently connected population of inhibitory neurons and excitatory neurons with both fast AMPA and slow NMDA synapses. In the present model multiple such populations are coupled together. The excitatory coupling *c*_*e*_ represents overlap between populations. The mutual inhibition, *c*_*ei*_, allows for competitive dynamics between populations, as does the value of *c*_*ie*_. Both of these connections provide similar dynamics to the results described in the next section (see, e.g., Table 1 in the weak coupling case). Thus, we have set *c*_*ie*_ = 0 for simplicity in order to study different inhibitory connectivity architectures, described below. We first consider two populations, connected as in Fig 1B, with the cross-excitation coupling, 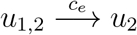, and 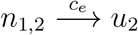, given as

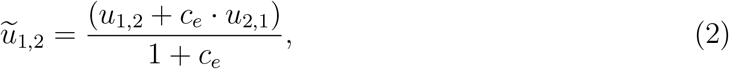

and similarly for 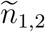 (i.e., let *u* ← *n* in Eq (2)). The mutual inhibition term for two populations, 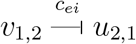, similarly is given as

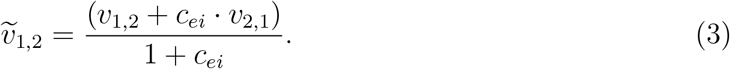

**Table 1.** Weak coupling summary of the dynamics for up to 4 active groups. The states are synchronous (S), anti-phase (A), nonsynchronous (N), clustered (C), symmetric cluster (CS), splay (L), and semi-splay (L3). By synchronous, we mean that all the oscillators fire together in-phase; anti-phase means the two oscillators fire a half cycle apart; clustered means that two groups of oscillators form that are synchronous within the group and out-of-phase between groups; symmetric clusters mean that there are equal numbers in each group; nonsynchronous means neither synchronous nor anti-phase; splay is the state: 0, 1/*N*, 2/*N*, …, (*N* − 1)/*N*; semi-splay is a state where 2 oscillators are synchronized and the other two are out of phase but not synchronized themselves. Thus, A, L, and N correspond to the out-of-phase (OP) oscillations described in the main text while C, CS, and L3 correspond to the mixed-phase (MP) oscillations.

| Active | EE | EI | IE | II |
| --- | --- | --- | --- | --- |
| 2 | S, A | S, A | S, A | N |
| 3 | S | S, L, C | S, L, C | C |
| 4 | S | S, CS, C, L3 | S, CS, C, L | C |

The denominators in both Eqs (2) and (3) allow for bounded excitation and inhibition, respectively. Increasing the number of coupled populations allows for different possible connectivity architectures, as shown in Fig 2. Using all-to-all connectivity (Fig 2A), the terms can easily be generalized from Eqs (2) and (3). For example,

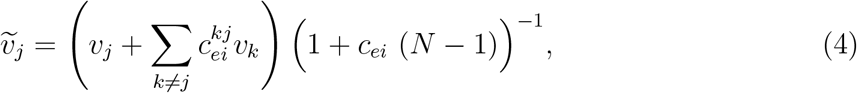

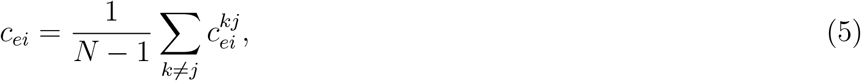

where 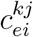 describes the coupling from the inhibitory component of population *j* to the excitatory component of population *k*. The cross-excitation equations are similar (i.e., let *v* ← *u* or *n*, *c*_*ei*_ ← *c*_*e*_, and 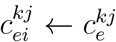 in Eqs (4) and (5)). The simplest and most tractable architecture to consider is uniform coupling. In this case the 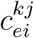 are constant, so that we relabel them by letting 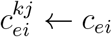, and the *c*_*ei*_ in Eq (4) is this same value (which, of course, is still equivalent to the formulation given in Eq (5)). The excitatory case is similar, and we impose the additional condition that *c*_*ei*_, *c*_*e*_ < 1 so that self-coupling is stronger than interpopulation coupling. We will refer to this setup as the uniform architecture (Fig 2B). We note that the uniform architecture is an ideal case, and we would expect the connection strengths to be at least somewhat heterogeneous. We may impose such a heterogeneity in different ways; for example, we may have inhibition that increases or decreases relative to the populations’ distances in either euclidean or feature space. One proposal for the observed capacity of working memory is interference of memoranda due to feature overwriting [33]. Stimuli with similar features may compete to be represented in working memory; for example, given two similar, sequentially-presented vibrotactile stimuli, the second may overwrite the memory of the first [34]. This motivates us to consider feature space and impose a distance-dependent coupling that decreases monotonically, as shown in Fig 2C for fixed *j*. We impose periodic boundary conditions, providing the network with a ring topology. Alternatively, we may draw each 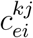 from a distribution (e.g., 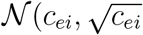), as illustrated in Fig 2D. Such heterogeneous architectures allow for additional dynamics, including differential quenching of synchronous populations and additional dynamic changes in binding.

**Fig 2.**
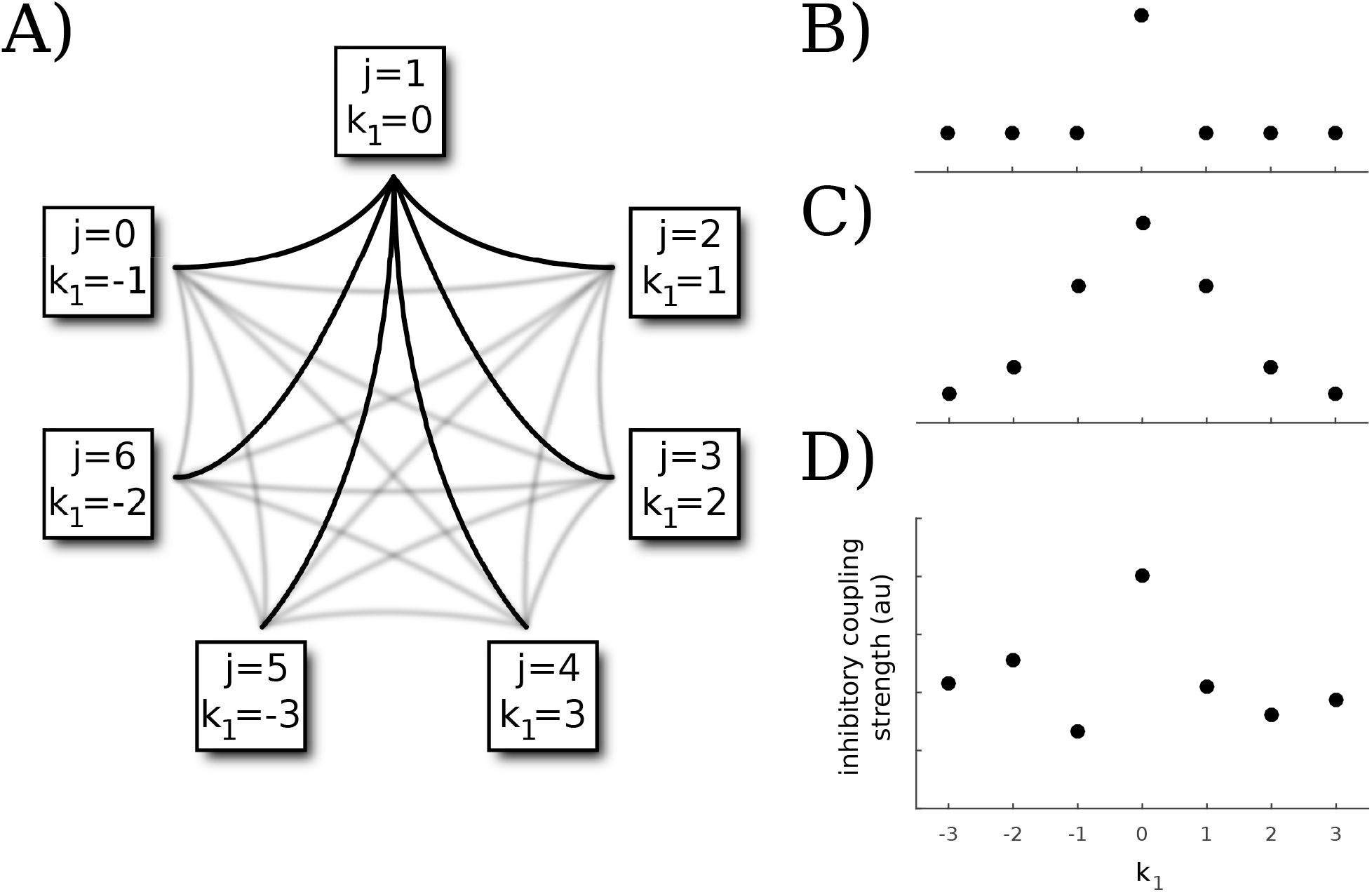
Connectivities for *N* populations. The networks of size *N* are coupled in an all-to-all fashion, as shown in (A) (here *N* = 7). Each network edge corresponds to bidirectional excitatory and inhibitory coupling with strengths *c*_*e*_ and *c*_*ei*_, respectively. We consider the inhibitory coupling strengths from population *j* + *k*_*j*_ (mod *N*) to population *j*, using the labeling scheme as shown in (A) (the populations are labeled from 0 to *N* – 1 for convenience with modulo arithmetic). Without loss of generality, we consider the connectivity of population 1 (*j* = 1) (black edges include population 1, gray edges do not). (B – D) Coupling values for *k*_1_ = 0 correspond to the self-inhibition strength *a*_*ei*_ for population 1, and those for *k*_1_ ≠ 0 correspond to *c*_*ei*_ values. We explored three different connectivity patterns for which we provide schematics: uniform (B), distance-dependent (C), and random (D). Note that for random connectivity, the graph may look different for each population, whereas the uniform and distance-dependent connectivities will produce identical graphs for all populations. In order to obtain winner-take-all dynamics, these inhibitory strengths are all positive.

Unless otherwise specified, the parameters used are *N* = 5, *τ*_i_ = 12, *τ*_*n*_ = 144, *C*_*e*_ = 0.001, *c*_*ei*_ = 0.03, *a*_*ee*_ = 14, *a*_*ei*_ = 10, *a*_*en*_ = 4, *θ*_*e*_ = 6, *a*_*ie*_ = 20, *a*_*ii*_ = 8, *a*_*in*_ = 0.1, *θ*_*i*_ = 5, *a*_*n*_ = 2, *β* = 1, *p* = 2. These parameters allow for tristability among three behaviors of interest: a low nonzero steady state, a low-amplitude oscillation around this low steady state, and a large-amplitude oscillation. In our model, “active populations” refer to populations that are engaging in large-amplitude oscillations. Note that we vary *N*, *τ*_*i*_, *τ*_*n*_, *c*_*e*_, and *c*_*ei*_ to study their effects below.

Our protocol for simulating working memory is to load a memorandum via a square wave pulse stimulus (*s*_*j*_ (*t*; *A*_*j*_, 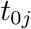, w_j_) in Eq (1)). Each stimulus is associated with the excitatory component of a specific population, so that a particular population may be selected for a memorandum. Multiple populations may be selected and provided stimuli either simultaneously or serially. Varying each of the amplitude *A_j_*, onset time 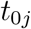, and width *w*_*j*_ may produce different activation patterns, so that the observed activity is a direct result of the selected parameters and sequence of feed-forward inputs.

Each stimulus corresponds to a feature of an item present in the environment, so that persistent activation of a population selected by a stimulus corresponds to that feature’s representation within the neuronal network. We allow for some overlap of excitatory populations, as represented by a nonzero *c*_*e*_ value. We then study how the system responds to patterns of stimuli, allowing their amplitudes, widths, and onset times to vary. We are particularly interested in phase synchrony and asynchrony of the large amplitude oscillations across the excitatory components of the populations. In our model, synchrony corresponds to different features that are associated with one another held active in working memory, while asynchrony corresponds to independent features held active. These behaviors correspond to binding and working memory capacity. We further study the existence and stability of these states while varying key parameters using XPP-AUTO.

However, since we obtain these network behaviors through a feedforward excitatory impulse to the excitatory components, we are further interested in what states are in fact accessible from some other given state. In systematically studying what patterns may be obtained from other patterns following a selective stimulus, the combinatorics involved quickly threaten to make the problem an intractable one. Thus, we limit our study to just two and three active populations, which we will refer to as diads and triads, respectively. We begin with either a diad or triad, and then use the protocol as described above, providing a single, selective stimulus of a chosen amplitude and width either to an active population or to an inactive one, and allow the network to evolve for some time afterwards. The resultant pattern is then determined to be accessible from the initial pattern. The outcome of stimulating the system is phase-dependent, as is the case in many oscillatory systems. Since we are interested in behaviors that may be more robust, we narrow our results to only include those that contain multiple-millisecond intervals of time that produce the same resultant activity for a stimulus of fixed width and amplitude. All of the included results may be obtained with a stimulus of fixed width and amplitude over intervals no smaller than 6ms. For simplicity, we only consider accessible operations for a network of uniform architecture with *N* = 5 when we explore the accessibility of operations involving diads and triads.

## 3 Results

The model produces persistent elevated firing states (above baseline levels) in response to inputs to selected populations in networks consisting of multiple populations, consistent with what has been observed in neurophysiological studies (e.g., [5]). This persistent working memory activation may occur either as steady-state or oscillatory firing rates, depending on the value of the relative speed of the inhibitory neurons. For the case of oscillatory dynamics, analogues of several critical features of working memory arise naturally and robustly, which, in addition to the persistent elevated activity, includes working memory capacity and binding. In contrast, for the steady-state case, it is difficult to obtain multiple populations active at once, and in the case of uniform connectivity, the activity would be indistinguishable (i.e., they would all be active with the same firing rate). An analysis and description of the mechanisms underlying the oscillatory behavior of an isolated *u* – *v* – *n* triplet is presented in Appendix A. Except where specified otherwise, the results detailed below are obtained using the uniform architecture. The same qualitative dynamics may be obtained with heterogeneous connectivities, including random and distance-dependent connectivities, under suitable bounds on the coupling coefficients 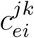.

Three basic types of oscillations may occur with coupled populations: out-of-phase (OP), synchronous (S), and mixed phase (MP). By OP, we mean that if there are *k* active populations, then each of the *k* occupy separate parts of a cycle. For example, if *k* = 2, then the two populations oscillate a half cycle apart, and if *k* = 3, each oscillate a third of a cycle apart. S means all populations fire in-phase with one another, and MP means that some populations are synchronized and others are out of phase. For example, it could be the case with three populations that two are synchronized while the third fires out of phase. When multiple populations are active in oscillatory states, they form a system of coupled oscillators. Hence, if the coupling between the populations is “weak”, it is possible to perform weak-coupling analysis on the system to gain insight into the possible attractors. In Appendix C, we perform this analysis, which shows that OP, S, and MP are all possible stable behaviors for 2, 3, or 4 populations. We note that although weak coupling analysis provides some insights into the mechanisms of the different states, the coupling we use in the full firing rate models is not weak, as evidenced by the ability of one population to suppress another.

These basic types of dynamics allow for individual memoranda to be represented in the case of OP oscillations, and for different binding scenarios to be represented in the case of S and MP activity. Greater complexity of memoranda and binding scenarios may be realized in the case of MP oscillations. The specific state of the network at any particular time, however, depends primarily on the particular pattern of inputs that have been selectively given to the excitatory components of the populations.

### 3.1 OP oscillations and distinct memoranda

OP oscillations of the interacting populations in the network (Fig 3) display the fundamental signature of working memory in the model. Each distinct memorandum being held active is associated with a different phase, similar to what has been proposed in previous work (e.g., [35], [27], [36], [37]). In contrast to some of this previous work, however, the present firing rate networks are self-contained. That is, they do not explicitly receive any external signal to organize their relative phase timings. Instead, mutual inhibition allows for competition among the populations so that only one population (or group of synchronously firing populations as discussed in section *3.3.1*) is active during any given portion of a cycle. Thus, for a given cycle duration (determined by conditions discussed in detail in sections *3.2* and *3.3.2*), there is a limit to the number of OP populations that the network may support. In the model this corresponds to working memory capacity. We will also refer to this capacity as the network capacity, as it is a fundamental property of the networked populations.

**Fig 3.**
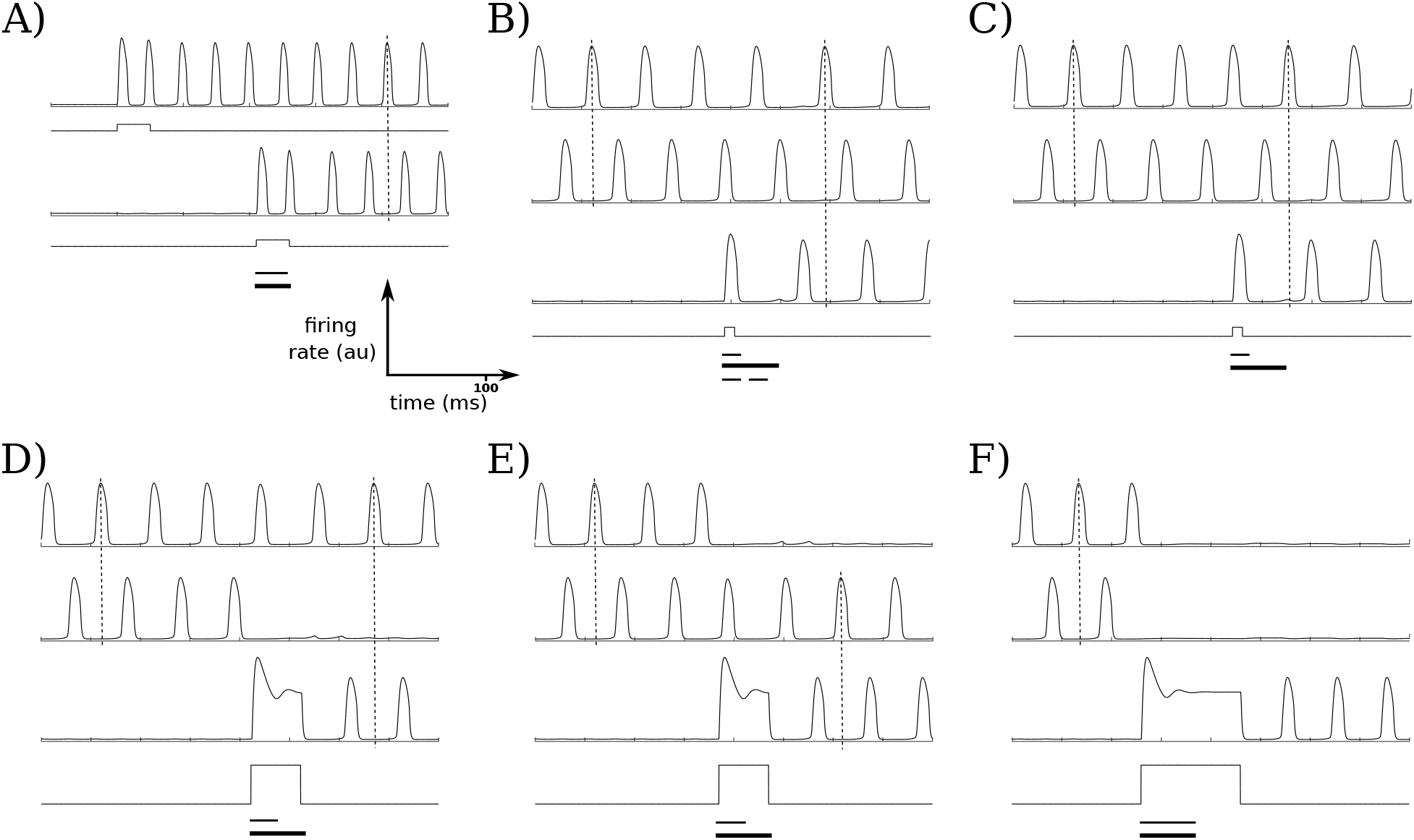
OP dynamics for up to 3 active populations. For a network size of 5 populations *(N* = 5) with uniform connectivity, all combinatorial possibilities for up to 3 active OP populations may be realized. Square waves in these and similar figures indicate the stimuli given to the population just above the given wave, and the vertical dashed lines in each plot allow for phase comparisons across different active populations. The first bar below each group of traces shows the interval of starting times in which the same stimulus (fixed amplitude and width) will produce the same result; the second bar shows the length of the period of the oscillation that is active before the stimulus is applied (note: the period may change after the stimulus is applied); the third bar in (B) is explained below. (A) The network starts at a nonzero, nonactive baseline firing rate. The first stimulus selectively activates the first population, while the other populations remain inactive with low firing rates. A second population is then activated; for these parameters and stimulus strength, almost any stimulus onset time will induce the OP state with 2 active populations, as the bars show. (B, C) Stimulating a third population with a short stimulus induces the OP state with 3 active populations. Either activation ordering may occur, depending on the phase the stimulus is presented; (B) and (C) s how the two different orderings. The third, dashed, bar below the stimulus trace in (B) s hows the interval of onset times that induce the OP state with three active populations; the first interval shows the onset times that produce (B) while the second interval shows the o nset times that produce (C). (D, E) Larger and wider stimuli may deactivate either of the active populations, so that the network remains in the OP state but with different active populations (WTS scenario). (F) Maintaining the amplitude of the WTS case but increasing the stimulus width allows the third selected population to deactivate both active populations and become the only active population (WTA scenario).

Depending on the timing and strength of the stimulus given to each population, a selected population may either oscillate out-of-phase with currently active populations in the network cycle, or it may compete with one or more populations, quenching their activity while remaining active itself. More concretely, consider the case of a 5-population network (*N* = 5) with a network capacity of 3 active OP populations. If no input is given, all of the populations fire at low, but non-zero rates. Selectively stimulating first one of the populations, and at a later time a second population, allows for these two populations to be simultaneously active, exhibiting OP oscillations as shown in Fig 3A. The result of subsequently selectively stimulating a third population depends on the strength, width, and onset time of the stimulus. Stimuli of sufficient width and amplitude can result in a winner-take-all (WTA) scenario (Fig 3F), where in the 3-population-capacity network the activation of the third population suppresses the first two populations sufficiently to become the only active population. Weaker stimuli allow for a winner-take-some (WTS) scenario (Fig 3D and 3E); in this case, the third population quenches one population (whose activity returns to baseline level) and becomes persistently active, leaving two OP populations. Both of these cases could represent selective forgetting due to interference. For example, they could correspond to situations in which attention is shifted to one item at the expense of other attended memoranda, effectively forgetting these items that had been held active in working memory.

If the stimulus is of moderate strength (that is, of sufficient amplitude and duration to activate the population, but not sufficient to quench another active population), then, depending on the onset time of the stimulus, the selected population may become interleaved with the other active populations in the current cycle (Fig 3B and 3C). In this case, all populations may fire OP, with the ordering of the newly activated population with respect to the already-active populations within the cycle determined by the stimulus onset time (e.g., Fig 3B vs 3C).

As each population becomes active, the oscillation frequency decreases, as illustrated in Fig 4. For any set of parameters, there is a limit to how many OP populations may be active, which we explore in greater detail below in section *3.2*. If the number of active OP populations remains constant, the frequency of oscillations also decreases with increasing network size. Thus, for example, if there are three active populations with OP dynamics, each population oscillates at just over 15 Hz for a network with *N* = 3, and at approximately 13 Hz for a network with *N* = 20. We see in Fig 4 that if either *N* or the number of active populations corresponding to distinct memoranda increases, the frequency of oscillations for any individual population trends downward towards the alpha band.

**Fig 4.**
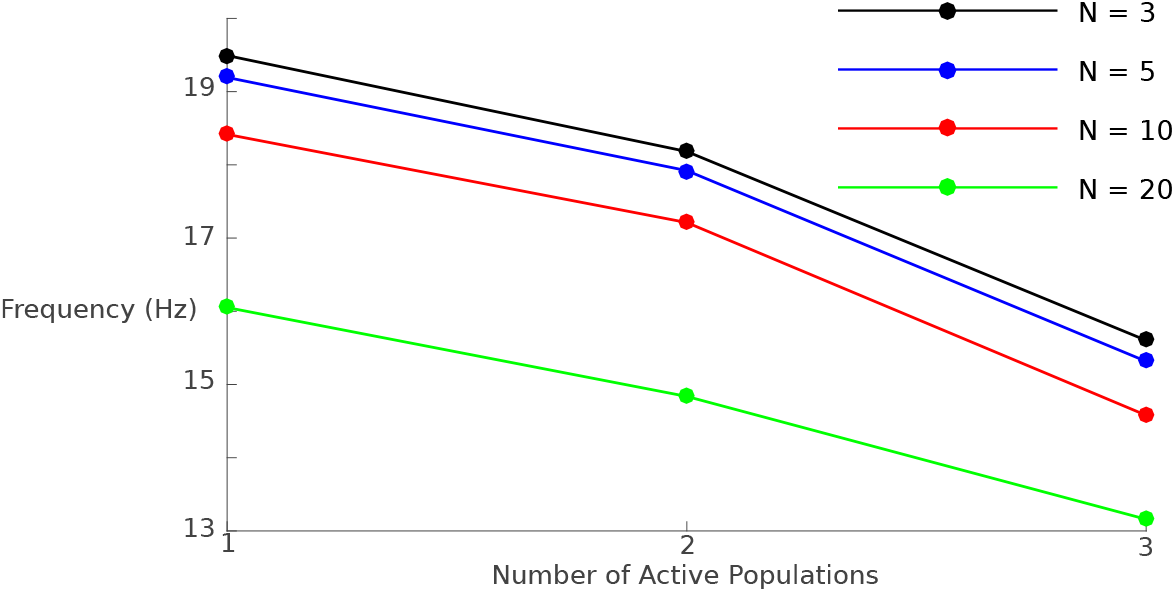
Frequency of population oscillation for different network sizes *N*. We examine the effect of increasing both system size and number of active OP populations on the oscillation frequency. In both cases, the frequency decreases monotonically. The frequency of interest may also be that based on the time between peaks of active populations (e.g., the time between the peak firing rates of population 1 and population 2 in a given network). This “interpopulation frequency” may be obtained by multiplying the given population by the number of active populations. For example, the interpopulation frequency for a network size of 20 when 2 populations are active would be approximately 30 Hz.

Since we are considering OP dynamics, we also note that the period between successive peaks in the case when, for example, three populations are active, is one third the overall period of the network. We may refer to the associated measure as the “interpopulation period/frequency”. As an example, let us consider a network size of *N* = 20 with three active OP populations. As mentioned above, the network oscillation, and thus the oscillation of any particular population, is 13 Hz. However, since three populations are active OP, the interpopulation frequency is 39 Hz. Depending on the distances between these active populations of neurons and the spatial resolution of the measurement in an experimental setting, then, increased activity near the alpha and/or gamma band may be detected.

### 3.2 OP oscillations and working memory capacity

As mentioned above, for any set of nontrivial parameters (e.g., *c*_*ei*_ ≥ *∊* < 0) there is a maximum number of OP populations that may be active, resulting in a finite network capacity. Both the mutual inhibition, *c*_*ei*_, and the timescales, *τ*_*i*_ and *τ*_*n*_, are critical in determining this capacity. We expect the value of *c*_*ei*_ to play a strong role in determining the network capacity since the OP dynamics fundamentally arise from mutual inhibition between the populations in the network (see sections *2* and *3.3.2*). The effect the timescales have on the capacity is somewhat more nuanced. The timescale of inhibition works differentially on the excitatory populations, depending on their phases. Briefly, we find that, in particular, larger values of *τ*_*i*_ increase the time the excitatory populations spend near zero more so than the time they spend away from zero, providing a greater window of opportunity for other populations to be active. We now look at this in greater detail.

We observe that there appear to be two qualitatively different features to the excitatory oscillation: (1) a pulse-like portion, where the activity rapidly increases and rapidly decreases, which we will call the “active phase”; (2) a portion where the firing rate stays very close to zero, which we will call the “quiescent phase”. Although the division between these phases is necessarily arbitrary, a natural threshold choice is the low fixed point, *u*_*_ (the baseline firing rate). Thus, for some fixed *j*, the active phase of a given oscillation, *T*_*a*_, is defined as the interval of time such that *u*_*j*_ > *u*_*_, while the quiescent phase of an oscillation, *T*_*q*_, is the interval of time such that *u*_*j*_ ≤ *u*_*_. Thus, *T*_*a*_ is a measure of the width of the pulse, while *T*_*q*_ is a measure of the time it spends near zero. Note that *T*_*a*_ + *T*_*q*_ = *T*, the period of the oscillation. *T*_*a*_ is mostly determined by the rise time of the inhibition, which is strongly affected by the excitatory population. In contrast, *T*_*q*_ is mostly determined by the decay time of the inhibition, which is only weakly affected by the excitatory population. See Appendix A for more detail. Thus, while both *T*_*a*_ and *T*_*q*_ increase as *τ*_*i*_ increases, we expect 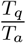 to increase as well. Indeed we find this to be the case; for example, with our original parameters (including *τ*_*i*_ = 12) and one active population, *T* ≈ 50.22ms, and 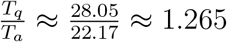. If we increase the timescale of inhibition up to *τ*_*i*_ = 20, *T* ≈ 76.35ms, and 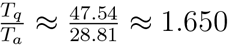. Thus, the ratio 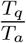 has increased by a factor of about 1.3. We therefore might intuit that more populations can be active OP as *τ*_*i*_ increases, since more pulses from other populations may “fit” between the pulses of an already-active population. That is, since there is a proportionally larger interval of time the population spends near zero than the interval of time it is active, there is greater opportunity for other populations to be active during this quiescent phase. We also note that while it is important for *τ*_*n*_ to be large enough so that the NMDA population may reactivate the excitatory population (see Appendix A), it otherwise does not affect the period of the oscillation very much. For example, with our original parameters (including *τ*_*n*_ = 144) and one active population, recall that *T* ≈ 50.22ms; increasing *τ*_*n*_ to 240 slightly *decreases* the period to *T* ≈ 48.94ms. (We note that it is not surprising for *τ*_*n*_ to have an inverse relationship with the period, since *τ*_*n*_ mostly controls the decay time of the NMDA; thus, *n* stays slightly higher for larger *τ*_*n*_, resulting in slightly faster activation times for *u* and *v*. Since *τ*_*n*_ has little effect on the decay time of inhibition, which is mostly determined by the time constant *τ*_*i*_, we see that increasing *τ*_*n*_ may in fact decrease the period.)

We may follow the oscillatory solutions in AUTO to determine exactly how the network capacity depends on the mutual inhibition and the system’s timescales. In the uniform case, the OP states are lost as folds of limit cycles or destabilize via torus or period-doubling bifurcations as *τ*_*i*_ increases or decreases with *τ*_*n*_ fixed (Fig 5A). By following these bifurcations, we may partition *τ*_*i*_ – *τ*_*n*_ space based on how many OP populations can be active for fixed *c*_*ei*_ (Fig 5B), or on the maximum number of OP populations that may be active for different values of *c*_*ei*_ (Fig 5C). In Fig 5 we see that we have open sets of *τ*_*i*_ and *τ*_*n*_ values that lie within physiological ranges where the network capacity is 3, matching experimental ranges of a capacity of 3 – 5 items in working memory. As we increase the number of populations, the capacity of the system remains more or less the same (Fig 5D), suggesting the capacity is only weakly dependent on the network size.

**Fig 5.**
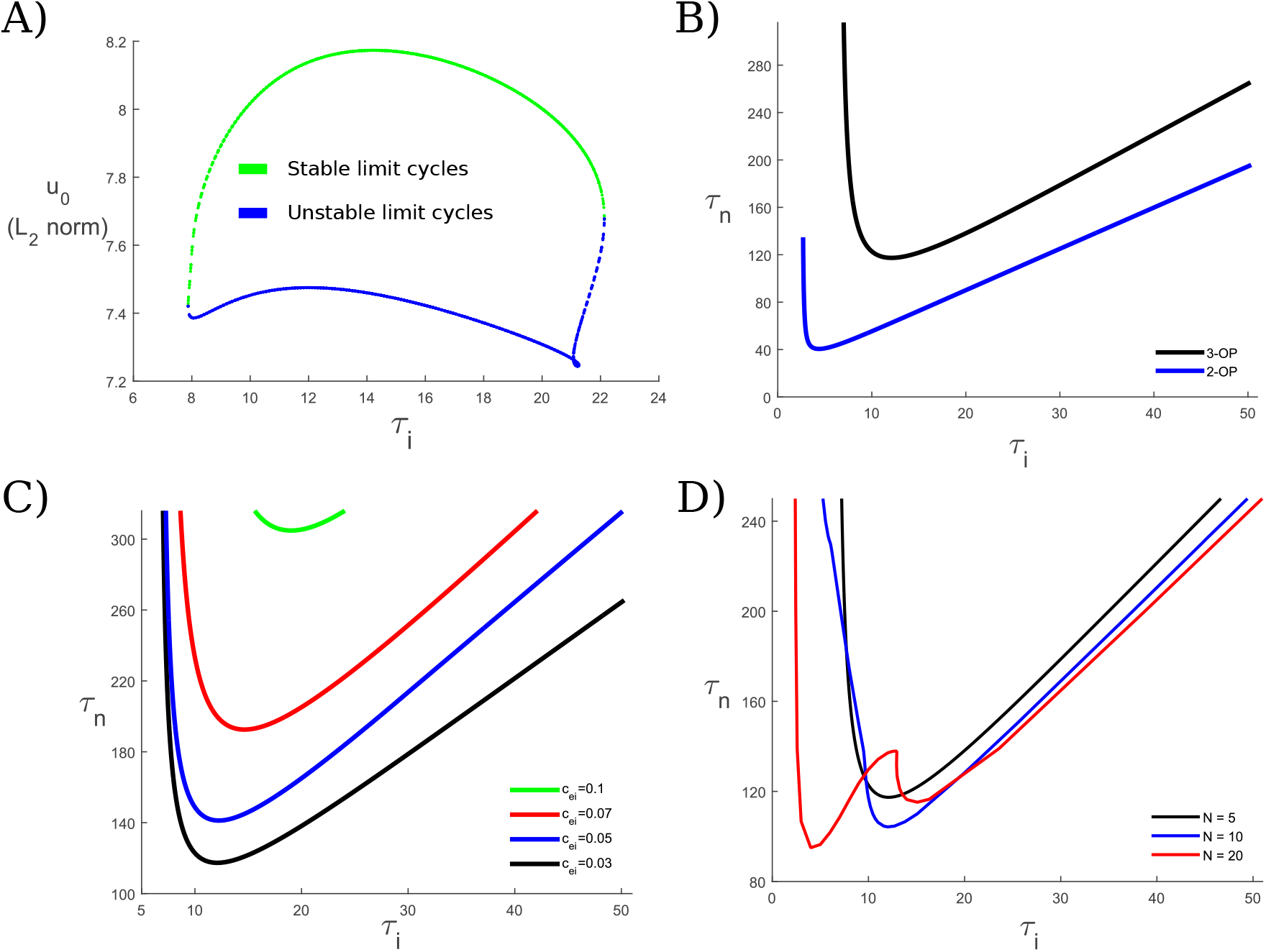
Stability of OP oscillations and working memory capacity. For fixed *τ*_*n*_, there is a window of *τ*_*i*_ values for which oscillations exist. We examine the existence and stability of the active out-of-phase populations, corresponding to distinct, single-featured memoranda. (A) Here, we look at the 3-OP state for a network of *N* = 5 populations, with *τ*_*n*_ fixed at 144 and the mutual inhibition *c*_*ei*_ fixed at 0.03. For *τ*_*i*_ too large or too small, the oscillations are lost as folds of limit cycles (for larger *N*, they may also be lost through torus or period-doubling bifurcations). (B) By following the limit points in (A) and keeping *N* fixed at 5, we may examine the dependence of the oscillatory states on both timescales, *τ*_*n*_ and *τ*_*i*_, for different OP states. Thus, we see how the capacity of the system depends on the timescales. Each curve is a curve of the limit points as shown in (A). Thus, the OP state with 2 active populations exists stably above the blue curve, and the 3-OP state exists stably above the black curve. (C) We may further examine how the capacity is affected by the mutual inhibition *c*_*ei*_. Here, the 3-OP state exists above each curve for different *c*_*ei*_ values as indicated. As the mutual inhibition increases, the minimum *τ*_*n*_ value that supports the 3-OP state increases. Thus, we would like to keep the mutual inhibition low enough to support the 3-OP state within physiologically realistic synaptic timescales, but high enough to allow for the WTA state. (D) If we fix *c*_*ei*_ = 0.03, we may further explore how the network size *N*affects the 3-OP state. Overall, as *N* increases the set of timescales that supports the 3-OP state does not change very much, generally increasing slightly.

In the case of the randomized architecture, the network may no longer have a capacity that is independent of which specific populations are active. As the interpopulation inhibition strengths 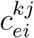 are sampled from a distribution (see Fig 2D and the discussion following Eq (5)), the interactions among populations will vary, so that the number of populations that may engage in OP oscillations can depend on which populations are stimulated. For example, suppose that N = 3, and take 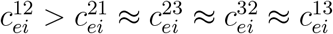. If the disparity between inhibition strengths 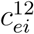 and 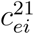 is sufficiently large, populations 1 and 2 may not be able to display OP activity, whereas populations 1 and 3 and populations 2 and 3 may be able to do so. Thus, there may no longer be a single network capacity, but rather a sequence of subnetwork capacities, depending on which populations are selected to be active. Exactly how much disparity among inhibition strengths may be tolerated while maintaining a given capacity is a nuanced question that may be of interest in future work.

### 3.3 S and MP oscillations and binding

Working memory is critical in binding. Here we describe how the patterns of synchronization that emerge from the model are rich enough to accommodate many aspects of the current understanding of binding from a working memory standpoint. The particular types of binding that may be obtained depend on the specific architecture in addition to patterns of input.

In all examples of binding, temporal synchronization of firing may play a significant role as the underlying mechanism. In working memory, activation of populations must be able to organize themselves to allow for the different dynamic and combinatorial types of synchronizations required for the different cases of binding. In the present model, we found a range of combinatorial arrangements and dynamics that emerge that are rich enough to map onto or underlie many of the aspects of binding.

#### 3.3.1 Binding vs. independent memoranda, and network capacity

All forms of binding may involve synchronous firing of different populations and circuits that are maintained active in working memory; bound features in our model therefore correspond to populations that display S activity. In contrast, the maintenance of different, distinct memoranda indicates persistent activation of multiple populations and circuits that do not fire synchronously, and so display OP dynamics. S and OP activity may occur together. When such MP (mixed-phase) activity occurs, the populations that oscillate synchronously correspond to different features of a memorandum, while populations that fire out-of-phase with one another correspond to independent components of multiple, distinct memoranda.

Thus, two capacities of the network may naturally arise: One associated with how many populations – or more accurately, how many groups of synchronized populations – may oscillate out-of-phase, and one associated with how many may oscillate in-phase. As we mentioned in section *3.2*, we refer to the first capacity as working memory capacity, or, due to its importance, the network capacity, and the second as binding capacity. We note that a working memory capacity is entailed either by nonzero coupling (i.e., one of *c*_*e*_, *c*_*ei*_, *c*_*ie*_ is positive), or by the temporal resolution of the network (i.e., how close oscillations may be before they are treated as synchronous). In either case, there is a minimum distance, say *η* > 0, that the excitatory peaks, e.g., may be to each other (in the first case, the minimum distance may arise due to the attracting or repelling basins of the synchronous state, for example). Thus, if *T* is the period of the network oscillation, the maximum working capacity would be 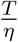, which is finite for any fixed set of parameters.

The binding capacity – i.e., how many populations may oscillate synchronously – may be larger than the working memory capacity. For example, for the uniform architecture with *N* = 5, we can have all 5 oscillate synchronously with the given set of parameters. Increasing the intrapopulation inhibition, *a*_*ei*_, decreases the binding capacity monotonically over a very narrow range: 5 populations may oscillate synchronously up to *a*_*ei*_ = 12.99, where the solution is lost at a branch point, while 2 may oscillate synchronously up to *a*_*ei*_ = 12.95. We note that the mutual inhibition, *c*_*ei*_, does not play the major role in limiting the binding capacity that it does in working memory capacity. For example, we can see from Eq (4) that if all *N* populations are active S in a given network, they oscillate as a single network with inhibition given by *a*_*ei*_.

Clearly, then, in the case of MP dynamics the total number of active populations may exceed the capacity of either working memory or binding. For example, the working memory capacity of the network shown in Fig 6 is 3. However, as we see in the case of MP dynamics (Fig 6E and 6F)), we may have 4 (or more) active populations. Whether or not we get S, OP, or MP dynamics depends on the stimuli parameters. For example, we see in Fig 6D that when a fourth population is stimulated while 3 OP populations are active, it may quench the activity of the third population and take its place, indicating that forgetting of the associated memorandum or feature has occurred, and features of 3 distinct memoranda remain (since 3 populations remain active OP). The fact that the stimulus was not particularly large suggests the network capacity is 3, which is indeed the case for these parameters. A new feature can also be added to an already existing memorandum, in which case the new feature is represented when a population synchronizes with an already-active population, either by the activation of a quiescent population (Fig 6C, 6E, and 6F), or else by the additional input to an already-active population (Fig 6D with the second stimulus). We note that the only differences between Fig 6D, 6E, and 6F are the onset times of the stimulus (the first stimulus in the case of Fig 6D) to the fourth population.

**Fig 6.**
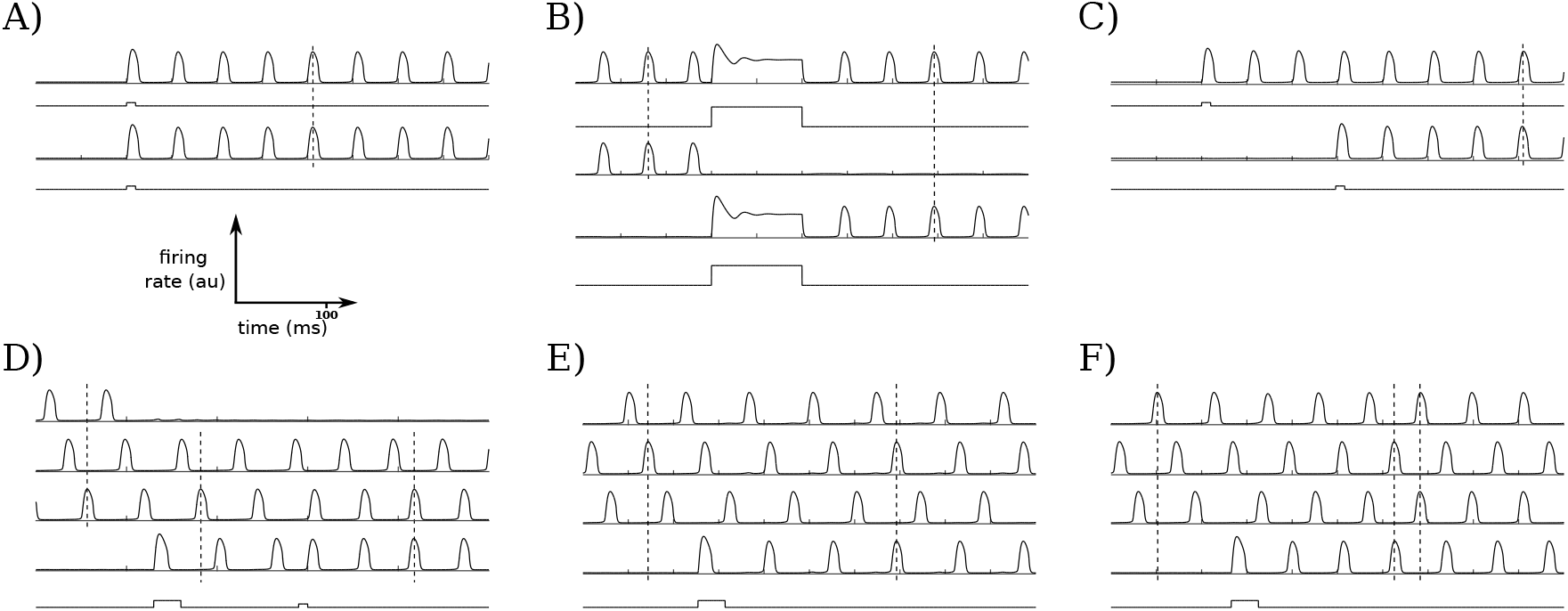
S and MP dynamics with simultaneous or sequential stimuli. (A – B) Simultaneous stimuli can cause the selected populations to exhibit S oscillations, whether they are activated from baseline as in (A), or while other populations are also active as in (B). (C) Sequential stimuli of the right timing may cause the selected populations to dynamically bind and oscillate S. (D) For networks that are already at capacity (3 OP populations in this case), a newly activated population may quench the activity of one population and continue oscillating OP with the remaining two. A subsequent stimulus to this active population may allow it to oscillate S with one of the two other active populations, and OP with the second. (E – F) Changing only the timing of the first stimulus from (D) may cause various dynamic bindings and MP dynamics. In (E), the network is still operating at capacity (3 OP groups), but there are more populations active (4 instead of 3). Similarly in (F), there are 4 populations active, but in only 2 OP groups, so that the network is still operating below its capacity.

#### 3.3.2 Two populations: Effects of different coupling strengths

Having seen that our model supports both OP and S dynamics that may map onto different working memory and binding scenarios, we look at the reduced case of two populations to distinguish how the interpopulation coupling affects the existence of these dynamics, as well as the case where only a single oscillator is active (SO). We note that the stable existence of the SO state is necessary for WTA dynamics; however, the SO state may also stably exist with lower coupling values that do not allow for WTA, and so it is not sufficient for WTA dynamics. The coupling strengths are crucial in being able to maintain any of these oscillatory dynamics. We see that in the coupling-strength space in Fig 7, only region *I* supports all three oscillatory states. Both OP (regions *I* and *II*) and SO (regions *I* and *IV*) states exist stably in bounded sets, so that if either *c*_*e*_ or *c*_*ei*_ values become too large, the states will be lost. In contrast, there is no indication that the S state only exists in a bounded region. If *c*_*ei*_ values are large enough with *c*_*e*_ small, the S state may lose stability (region V). However, we see from Eq (2) that as *c*_*e*_ → ∞, each excitatory population only gets excitation from the complementary population, so we expect synchrony to continue to stably exist for arbitrarily large *c*_*e*_ values. All three oscillatory states appear to exist stably when either *c*_*e*_ or *c*_*ei*_ is zero as the other value approaches zero (i.e., along one of the axes near the origin), suggesting that weak coupling is sufficient for these states to exist. In Appendix C, we explore the attracting states in the weak coupling limit.

**Fig 7.**
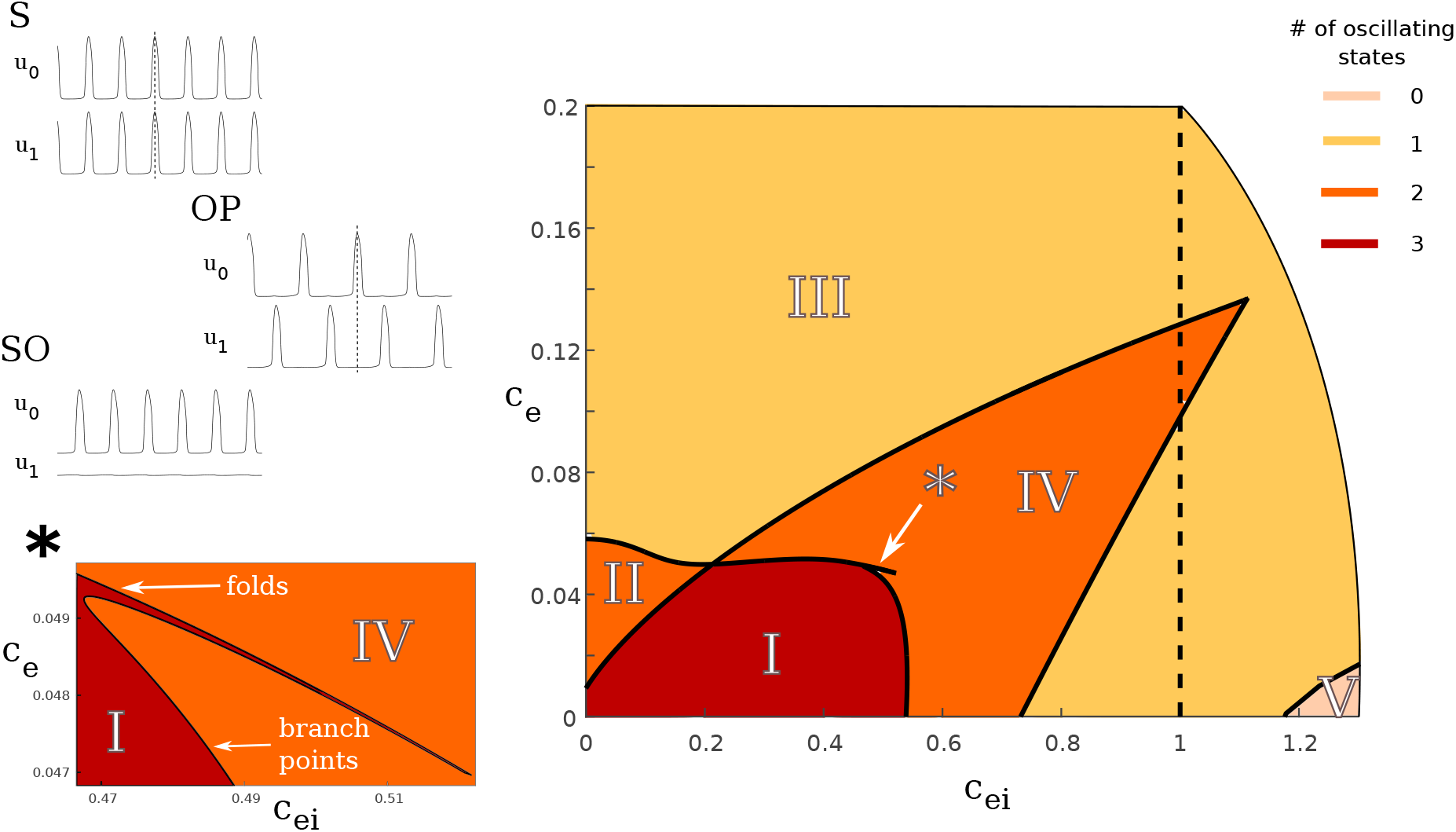
Stability as a function of coupling strengths for *N* = 2. We examine how the existence of stable states in the two population case depend on the coupling parameters *c*_*e*_ and *c*_*ei*_. The three states we examine are S, OP, and SO (the left panel shows the corresponding time traces). The colors indicate how many of these states are stable for the given coupling values. We have restricted the coupling values to be no greater than 1, so that interpopulation coupling is no stronger than intrapopulation coupling. The states that exist stably in each region are: (I) S, OP, SO; *(II)* S, OP; *(III)* S; *(IV)* S, SO; (V) no oscillating states. The dashed line at *c*_*ei*_ = 1 indicates the interpo pul ation and intrapopulation inhibition are the same. Thus, all of the dynamics of interest exist stably when the interpopulation coupling is less than the intrapopulation coupling, as desired. The inset (*) shows the cusp bifurcation as indicated in the right plot, where folds of limit cycles and branches of limit cycles come together to form the upper right portion of r egion (I).

Although the general features in Fig 7 match our intuitions, some of the particular features are unforeseen and may warrant further scrutiny. We now look more carefully at the bifurcations that define the regions that the S, OP, and SO states stably exist in coupling-strength space. The S state is only demarcated by a curve of branch points at large *c*_*ei*_ values (left curve of region *V*), and so exists for all relevant coupling strengths (*c*_*e*_, *c*_*ei*_ < 1). Remarkably, it is stable when *c*_*e*_ = 0, even for *c*_*ei*_ values far from 0. The region of definition of the OP state is more complicated. It is defined by a curve of folds of limit cycles (top curve in regions *I* and *II*) that meets in a cusp with a curve of branch points (right curve in region *I*). These two curves, whose juncture is shown in greater detail in the inset in Fig 7*\**, are not monotonic. We see that that there is a small interval of *c*_*e*_ values around 0.06 where the OP state exists stably for small *c*_*ei*_ values, but is lost for higher values (e.g., *c*_*ei*_ not quite 0.2). Surprisingly, once the OP state is lost here, we initially can only get the S state. As *c*_*ei*_ increases further, we may also attain the SO state. Perhaps the most unanticipated feature of the OP region occurs near the local minimum of the curve of folds of limit cycles (again, the top curve in regions *I* and *II*). There is a very small *c*_*e*_ interval, near *c*_*e*_ = 0.05, where the the OP state is lost and then regained as *c*_*ei*_ increases, before finally being lost again for large *c*_*ei*_. The region that defines the SO state does not hold much surprise. It is defined by a curve of folds of limit cycles (top curve in regions *I* and *IV*) that meets in a cusp with another curve of folds of limit cycles (right curve in region *IV*) that arises from a subcritical Hopf bifurcation. Thus, as *c*_*ei*_ increases, the SO state is lost to a high fixed point, so that the system may display steady state bistability between a down state and an up state with the same inhibitory timescale as for the case of oscillations. However, a much larger input is required to switch between active populations in this case.

For example, suppose *c*_*e*_ = 0.001 and population 1 is active. If *c*_*ei*_ = 0.7 (“after” the Hopf, so that population 1 has a large-amplitude oscillation), stimulating population 2 with a width of 50ms and an amplitude of 3 (arbitrary units) is sufficient to allow population 2 to become active and quench the activity of population 1. However, if *c*_*ei*_ = 0.8 (“before” the Hopf, so that population 1 is in the up state), a stimulus to population 2 with the same width requires an amplitude of greater than 36 in order for population 2 to activate, sending population 1 back to the down state. This up state is only stable for smaller *c*_*e*_ values; for larger *c*_*e*_ the only stable state of the system is S activity.

#### 3.3.3 Feature binding

The dynamics that emerge within the working memory networks are sufficiently complex to map onto and potentially underlie many phenomena, including static spatiotemporal memoranda, dynamic binding (described in the following paragraph), and illusions (discussed in section *4*) or forgetting that result from “overloading” the system. The model allows for mixtures and changes in synchronous oscillatory dynamics that can operate as dynamic changes in bound features, in addition to changes of focus.

Dynamic binding is required in working memory in order to quickly associate and dissociate different elements together [33], such as different features of an item in the environment. In the present model, this means that items must be able to be both synchronized and desynchronized in response to different stimuli. We have seen above in Fig 6 how such synchronization may occur via either simultaneous (Fig 6A and 6B) or sequential (Fig 6C, 6D, 6E, and 6F) selective stimuli. Such synchronization allows different features to bind together into one memorandum. By allowing for heterogeneous coupling, an activated population may have differential effects on other populations. For example, by employing a distance-dependent connectivity (as described in section *2*), activating a population can desynchronize already-active S populations. In Fig 8B, two S populations are firing at gamma frequencies. Attentional mechanisms or environmental changes (e.g., a car begins to move) may cause the synchronized binding to dissociate, with the features of individual objects changing or becoming independent objects in working memory, while the net background continues to be maintained (but perhaps out of “focus”), with a concomitant shift in frequency for the objects of attention to the alpha range. This is consistent with neurophysiological experiments that have reported increased alpha frequency and shifts from gamma activity with increasing working memory load [6, 24, 25]. In addition, increased alpha-band oscillations have been reported to play an important role in mental processes related to attention and memory [38–41]. Other dynamic changes in the features of objects could also potentially be accommodated, such as that illustrated in Fig 8A. An object is initially held in working memory with its features synchronized (e.g., a red traffic light). A new input occurs (e.g., the light turns green), and the stoplight is then stored in working memory with the new feature (green traffic light vs. red traffic light). Thus, the present model accommodates both binding and unbinding upon the activation of a quiescent population.

**Fig 8.**
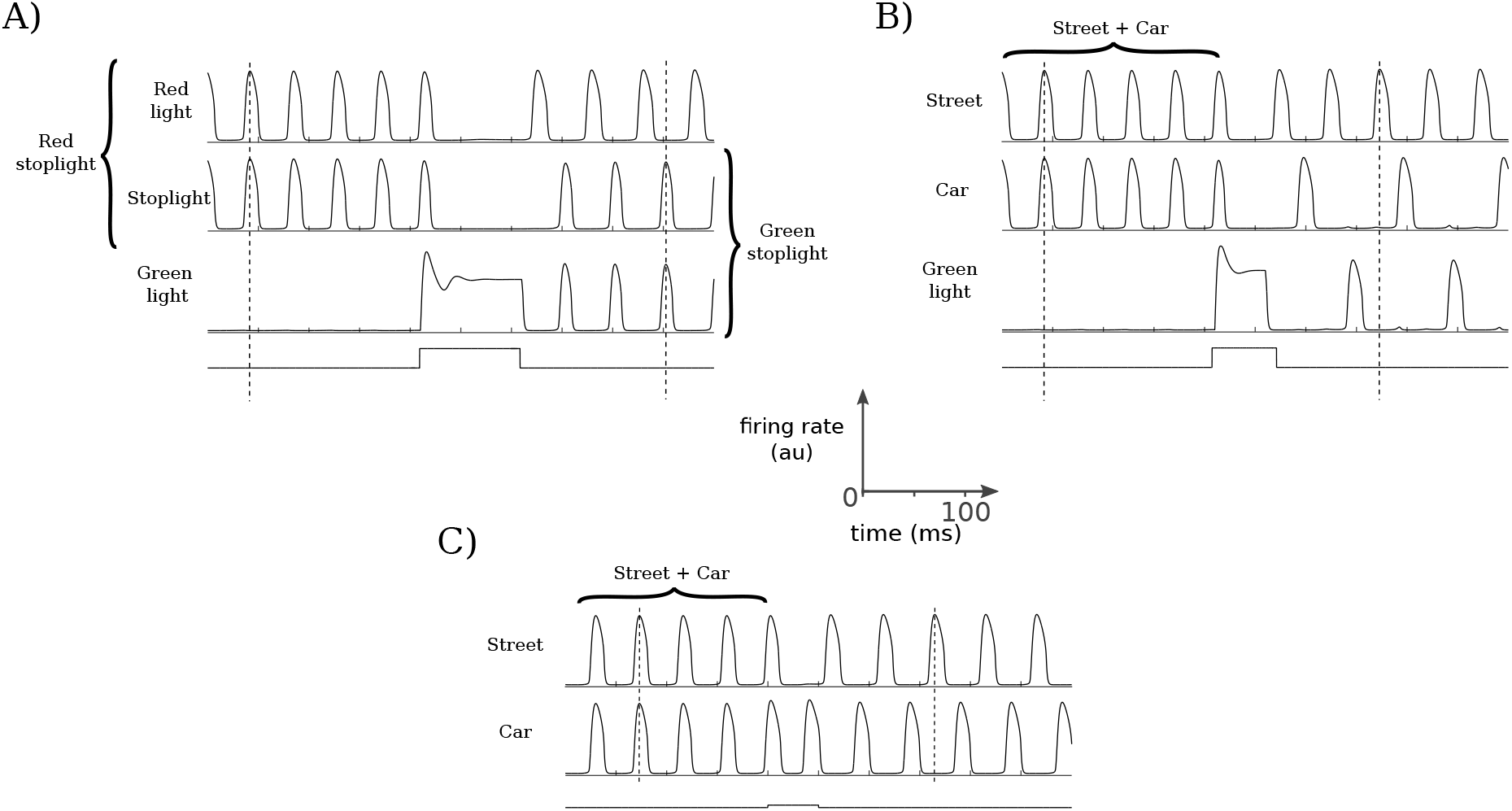
Feature binding examples. For simplicity, memoranda that may require several different bound features are represented as a single oscillating population (e.g., a street). We note that an asymmetric architecture is required to realize the dynamics shown in (A) and (B), as otherwise the newly stimulated population will affect the already-active populations symmetrically. Here, distance-dependent connectivity is used, along with the usual network size *N* = 5. (A) Feature binding can be achieved through the synchronous firing of the individual features as in the left half of (A). A new input causes an unbinding and a new binding of an object with a modified feature. For example, the initial binding might represent a *red stoplight.* The light changes to green, causing the *stoplight* to bind with *green light*, while the memory trace of the *red light* still persists (and perhaps is required for comparison purposes for subsequent action – i.e., to go). (B) Feature binding in which a new input (i.e., focus) results in the decoupling of a single memorandum (e.g., some background such as a *street* and stationary *car*). The change of a *red light* to a *green light* causes the *car* to move and decouple from the background so that there are two separate memoranda now, while the trace of the background (diminished in detail) maintains itself. Note that a shift in the distribution of frequencies present occurs as a result of the relative synchronizations and desynchronizations. (C) Here, the *green light* is not observed; rather, the *car* is simply observed to begin moving, and so is now perceived as a separate object from the *street*.

#### 3.3.4 Variable binding

The combinatorial dynamics emerging within the model are rich enough to produce variable binding as is characteristic, for example, of language and abstract reasoning within the context of working memory. Below, we illustrate some simple examples from the model consistent with the most expected and widespread approaches to variable binding and language (e.g., the SHRUTI inference network [42]). These utilize temporal phase binding and represent simple predicate calculus rules. Consider the predicate calculus rule

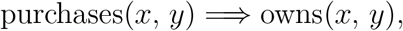

where *x* and *y* represent variables in the above rule, and a Boolean query (for example, as presented in Feldman [12]), such as “Does Tom (*x*) own a book (*y*)”. In most common inference network models, for example, SHRUTI [43], separate clock phases are assigned to the pairings of *Tom* with *owns* and *book* with *owns*. *Tom* here represents a variable agent who could come to own something (a *book*, for example) through *purchases*. There are other possibilities, of course, and they could all be linked or synchronized with the *owns* relation. We note that while a mediator circuit is implicated in the SHRUTI inference network, here the relevant association may occur in a single step as a result of synchronous firing due to temporal associations and the pattern of input, as shown in Fig 9. The active populations in working memory could represent an *owns* node (e.g., the active population in line 1 of Fig 9A), and *Tom* (as an agent node; e.g., the active populations in lines 2 and 3). The active populations of *Tom* and *book* then become associated and paired as a result of the activation of a node or population representing *purchases* in working memory (active population in line 4 of Fig 9A). Then, over several clock cycles “Tom owns a book” becomes established, or possibly instantly as here with the activation of the *purchases* node (i.e., population). Additional populations could become activated and linked via synchronization, such as “Tom purchases *The Awakening*” where *The Awakening* would become synchronized with a *book* node. We note that, whereas *Tom* requires two populations to be active in this example in order to form distinct bindings with both *purchases* and *owns*, different, *n* : *m* locking ratios or aperiodic oscillatory dynamics could allow for such unambiguous bindings while just one population is maintained active for *Tom*, as we touch on at the end of section *4*.

**Fig 9.**
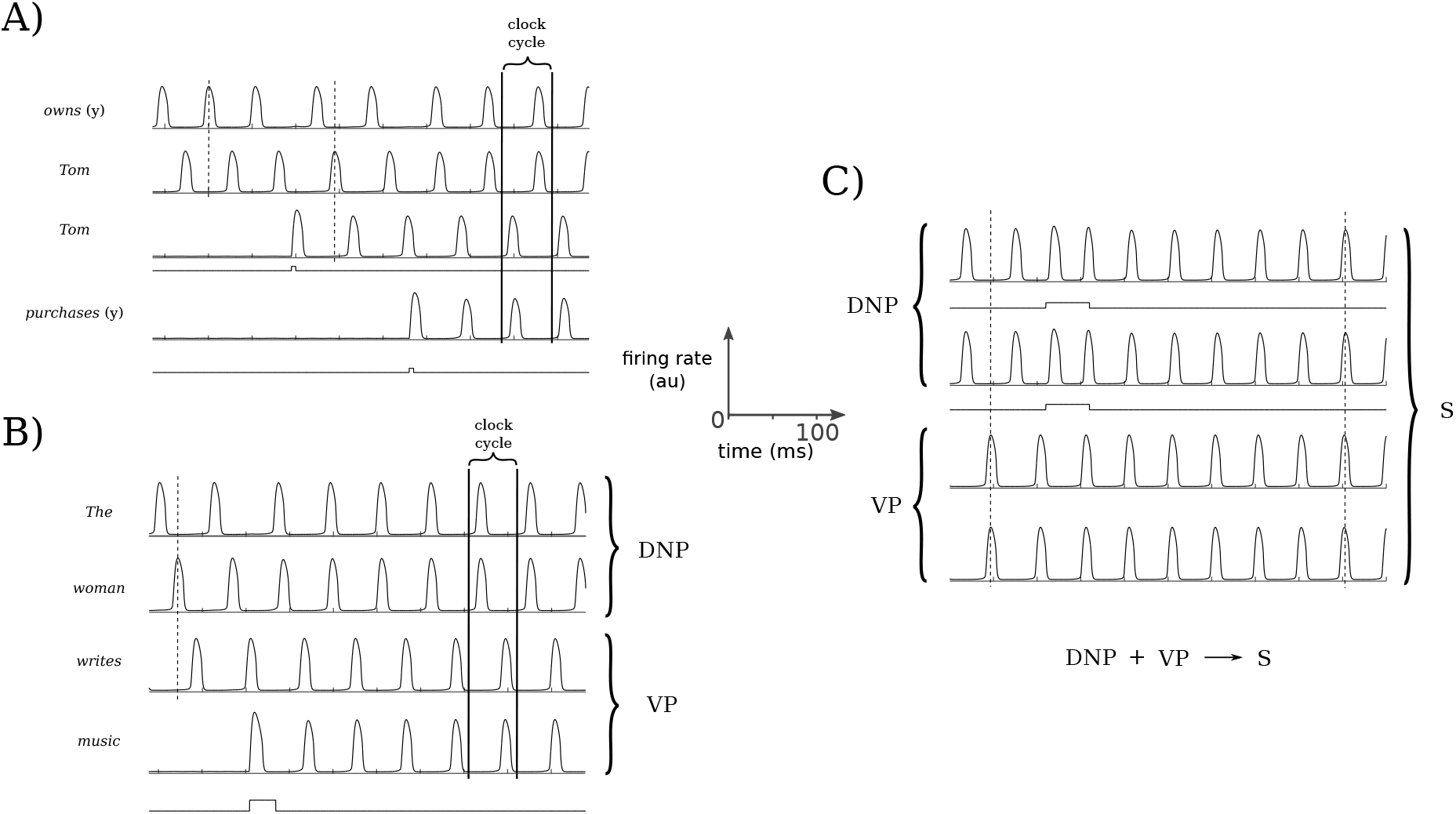
Variable binding, simple predicate caclulus, and sentence construction examples. Specific sequential inputs result in a cascade of bindings that emerge, following one another within a single clock cycle. (A) A fragment of the working memory system demonstrating how an inference emerges and is established in working memory. Here, the predicate calculus rule examined is purchases(*x*, *y*) ⇒ owns(*x*, *y*) (from an example in Feldman [12]). After a query is made (“Does Tom own *The Awakening?”*), a statement is provided, “Tom purchased *The Awakening”*. This statement first activates a second instantiation of *Tom*, and then the verb *purchase s(y),* causing *owns(y)* and *Tom* to synchronize so that the inference is made. (B) An illustration of that same combinatorial structure is shown as it could apply towards a mechanistic realization of a phrase structure grammar. This could be based upon an already established or innate structure in the cortex. Upon reading the sentence, the words are sequentially input and the appropriate nodes are activated and bound in working memory, forming a determinate noun phrase (DNP) and a verb phrase (VP). For clarity, bindings between nodes and variables (i.e., words here) are not explicitly illustrated. (C) A final binding occurs, as the components of the DNP receive selective and equal simultaneous stimuli, binding all of the components together to form a sentence (S).

We can also consider how the combinatorics allowed through the present model’s dynamics may apply in language by illustrating how they may facilitate the presence of grammars. We note that the predicate-argument structure used above (e.g., owns(*x*, *y*)) fits well within some dependency grammars. However, the model is agnostic to any particular type of grammar, and we now consider an example as might be implemented within a phrase structure grammar, illustrated in Fig 9B and 9C. The words are represented by the activation of populations (presumably stored in the structure of networks in long-term memory) and become synchronized with the appropriate activated nodes. For clarity, we have simplified the example so that the nodes, and therefore the bindings of words to nodes, are not displayed. For example, a determiner node is first activated and then bound to *the*, but in Fig 9B we only show the activity of the population corresponding to *the*. A noun node is then activated, which is bound to the variable *woman*. A verb node is activated next, which is bound to *writes*. Finally a second noun node is activated and then bound to *music*. This results in the binding of the determiner node and noun node of the subject into a determinate noun phrase (displaying the phrase structure rule D + N → DNP), and within a single clock cycle the dynamic binding of the verb with the direct object to form a verb phrase (V + N ? VP). These states are activated sequentially within a single clock cycle, and thus could be recognized as a grammatical sentence maintained in working memory based on the phrase structure rule that a DNP followed by a VP produces a grammatical sentence S. Alternatively, the sequential DNP and VP representations could be input with a subsequent final synchronization of the DNP and VP taking place to produce S, and recognized as a grammatical sentence that is maintained in working memory, as illustrated in Fig 9C.

Ungrammatical sentences could be recognized when bindings occur that do not correspond to valid production rules. We do not consider here the specific mechanism or details by which the particular variables become bound, but rather show how they can dynamically emerge within working memory representations. In principle this could arise via some mapping and closeness of the associations in that cortical map, “hardwired” architectural associations in long-term memory, or some combination.

### 3.4 Accessible operations

We have shown that the dynamics of our model map onto a number of working memory and binding examples. Indeed, it appears that the relevant attractor patterns are exactly the combinatorial possibilities, limited by the network size and the number of MP patterns available. Beyond ascertaining what states are available to the network, we are further interested in how the network can transition from one pattern to another. For example, in Fig 3B we see that if a third, inactive population is stimulated, the network can transition from 2 OP populations to 3 OP populations. We will consider this to be an operation available to the network; i.e., the operation of transitioning from 2 OP populations to 3 OP populations by providing a stimulus of the right strength and timing (see section *2* for further protocol detail). Different patterns may require a different number of operations to attain from a given starting pattern. We refer to a transition that requires *n* selective, sequential stimuli to instantiate as an *n* th-order operation. We may ascertain the capabilities, and thus to some degree the plausibility, of the model more systematically by delimiting the available operations. Determining whether a pattern is accessible through a first-order or a higher-order operation may also allow for predictions to be made from the model, such as the timings involved in certain cognitive processes, based on whether a particular pattern is accessible from another one directly with a 1st-order operation, or whether a 2nd- or higher-order operation is required. We restrict the number of active populations to either two (diads) or three (triads). Our parameter choices are as above, so that WTA is always an accessible state. Thus, if we begin with a diad or triad, the only resultant patterns of interest are also diads and triads.

Fig 10 shows what operations related to diads and triads are available to the network, under the limitations described above and in section *2*. In each gray circle, 1st-order operations are displayed; we may determine available higher-order operations by stringing together sequences of 1st-order operations. For example, we may transition from an OP diad to an S triad as a 2nd-order operation by stimulating an inactive population so that the network first transitions to an OP triad, and then stimulating an active population so that the network then transitions to an S triad. In particular, examining Fig 10B and 10C reveals that the network may transition from any diad or triad to any other (through transitions involving only other diads and triads) using no higher than a 3rd-order operation (i.e., no more than 3 selective, sequential excitatory stimuli are required).

**Fig 10.**
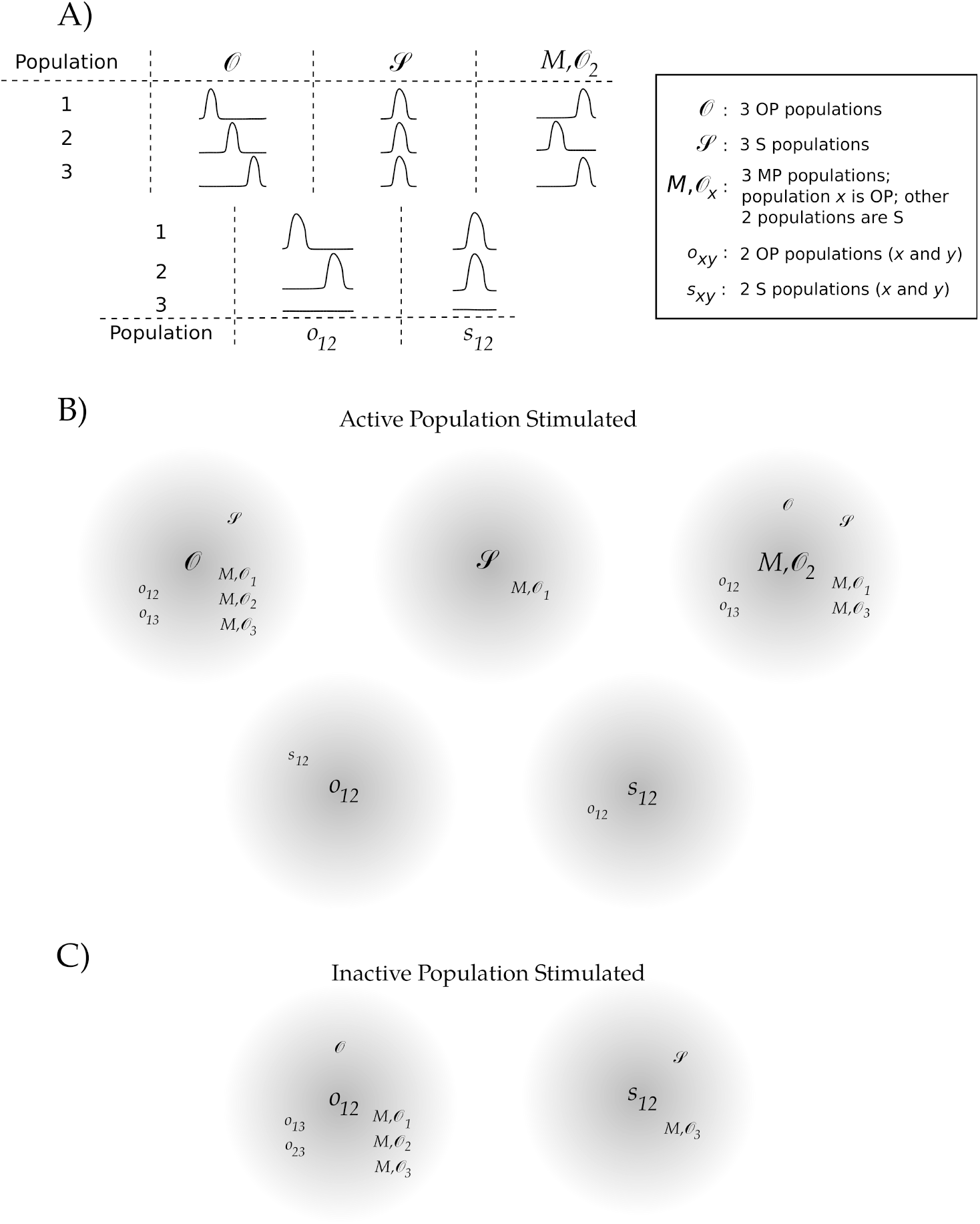
Accessible diad and triad operations. (A) Using *N* = 5, a stimulus is given to each of the activity patterns shown. For (B) and (C), the starting activity pattern is indicated in the center of the gray circles, while the subscripts indicate which population was stimulated. The observed resultant activity patterns are indicated in smaller text around the circumference of the gray circles. For (B), one of the already-active populations, population 1, was stimulated, whereas an inactive population, population 3, was stimulated for (C).

Since the stimulus is excitatory, the stimulated population will remain active. Thus, every beginning pattern will have at least two resultant patterns that will not be accessible for a given selected population that receives a stimulus (e.g., beginning with *O* and stimulating population 1 will ensure that *s*_23_ and *o*_23_ are not accessible – see Fig 10A for a notation key). By changing the population that receives the stimulus, other patterns become available in a way that is obvious by inspecting Fig 10. For example, we see in Fig 10B that an MP triad and OP diads and triads can transition to most of the remaining diads and triads: Discounting order, there are 5 possible triads and 6 possible diads; disregarding which population receives the stimulus, we see that both an MP and an OP triad can directly transition to 6 of the 10 patterns that are different from themselves. By contrast, the S populations have fewer populations accessible by 1st-order operations. This is in part an outcome of maintaining a uniform connectivity. In order to differentially affect two S populations, one of them must receive a stimulus, as any other population will affect the two S populations symmetrically. So, for example, it is as expected that the only accessible patterns for 3 S populations are the ones consisting of 3 MP populations where the selectively stimulated population oscillates out-of-phase with the remaining two. The results for *s*_12_ as shown are likewise entirely expected.

One of the patterns that is difficult to directly obtain from a diad or triad is an S diad (e.g., *s*_12_). We were only able to obtain this pattern by stimulating one of the active populations in an OP diad (e.g., *o*_12_). Thus, going from any other pattern to an S diad requires at least a second order operation. For example, transitioning from an S triad to an S diad is at least a third order operation: with the first stimulus the network may transition to an MP triad, with the second the network may transition to an OP diad, and with the third the network may finally transition to an S diad. We note that there may be multiple paths that allow one pattern to evolve to another pattern. For example, an alternative route from the S triad to diad would be through a WTA scenario: a strong first stimulus can cause the selected population to quench the activity of the other populations, and a second stimulus to a nonactive population could cause the network to transition to an S diad.

These results show that there are a number of options to get from one pattern to another for two and three active populations, even with strong limitations on the network architecture and the stimulus protocol. It may be of interest in future work to loosen some of the above restrictions (for example, examining heterogeneous networks or having the stimulus also drive the inhibitory components), and to further quantify the level of difficulty for a particular transition (e.g., some transitions are much less dependent on the stimulus parameters than others).

## 4 Discussion

This work presents a physiologically based firing rate model that produces working memory behavior and dynamics as observed in neurophysiological experiments, insofar as selected populations can exhibit persistent activation by maintaining above-baseline firing rates in response to specific inputs. In particular, we have shown that oscillatory dynamics in the network’s persistent states can provide a basic mechanism for, play a critical role in, and facilitate the establishment of many fundamental properties and aspects of working memory function. Specifically, by adding phase information to active working memory populations, oscillatory dynamics provide a simple, but perhaps crucial, ingredient to allow for physical or abstract items to be represented as either distinct or bound. These oscillations may also play a fundamental role in facilitating the stable persistent activation of working memory networks while enabling rapid transitions between different active network states, as is critical in cognition in general and in working memory in particular, including in a range of binding scenarios maintained in working memory. We have shown that the oscillatory attracting states that emerge naturally using even a simple uniform architecture provide a means for these rapid transitions between different stably bound or unbound configurations, allowing inputs and memoranda to be dynamically and combinatorially coupled and decoupled. Altogether, this suggests that the capacity observed in working memory may be the result of optimizing the potential competing requirements of distinct vs. bound information and maintaining stable activations vs. allowing for rapid transitions in those activations.

The oscillatory dynamics are robust across a wide range of biologically reasonable parameter values and emerge at the level of a single population. However, as indicated above, richer and more complex behaviors may be accessed with networks consisting of increasing numbers of populations. The possible behaviors appear as bifurcations in various critical network parameters. For example, stable OP oscillations appear as the mutual inhibition, *c*_*ei*_, increases. Such oscillations correspond to a nontrivial working memory capacity in the model as each memorandum may be distinguished based on its firing phase. However, these oscillations or their stabilities may be lost as *c*_*ei*_ increases further, at, for example, folds of limit cycles or torus bifurcations. Period doublings may occur as well, potentially allowing for more complex frequency-couplings among the active populations. Thus, in general, mutual inhibition both allows for a non-trivial working memory capacity and simultaneously limits that capacity. Stable S and MP oscillations occur as the interpopulation excitation, *c*_*e*_, increases. However, for *c*_*e*_ too large, the network is only attracted to a bulk oscillatory state. While synchrony is desirable in some binding situations, too much synchrony can be pathological, so it is necessary to maintain low-level excitatory connections between distinct populations.

The network is also capable of producing steady-state firing dynamics under certain conditions and parameter values (e.g., lower GABA time constants). However, we have found that, in comparison to such steady-state dynamics, oscillatory dynamics allow for much richer behaviors, including the maintenance of multiple, distinct, multi-featured memoranda, and rapid transitions in sequences of activations. Multiple distinct items may be maintained because multiple populations or memoranda may occupy different phases within the overall “carrier” cycle. The given capacity of the system varies with the network’s size and other parameters. However, we found in the present study that 3 simultaneously OP active populations may be maintained across a broad range of parameters (with or without binding), consistent with the 3 – 5 item working memory capacity that has been reported across different modalities [7], including for visual and verbal tasks [7, 12, 44]. The model provides a natural explanation for the capacity of working memory, apart from the system’s bifurcations discussed above, in that the excitatory coupling *c*_*e*_ *c*can greatly increase the basin of attraction of synchronous states. Thus, two populations may only get so close before they will tend towards synchrony, implying there may only be a finite number of OP populations.

The basic architecture of the model was designed with short time constant receptors (AMPA and GABA) and longer time constant NMDA receptors, consistent with circuitry found in areas of the cortex and associated in working memory with gamma-band oscillations [45]. We note that gamma and theta frequencies have been attributed to various receptor and molecular messaging time constants (see, e.g., [42, 46]), and the present model is consistent with such explanations, while additionally suggesting that network-level dynamics may also contribute to neuronal activity observed in different frequency bands. In the present model, oscillations occurred with frequencies that could be measured as ranging from alpha to gamma bands. For example, for most parameter values, the peak-to-peak period of oscillations within a given working memory cycle corresponds to either gamma or beta band oscillations. As multiple populations or memoranda become active within a given working memory cycle, however, while the overall peak-to-peak period remains in the higher frequency bands (gamma or high beta) in a working memory cycle, the peak-to-peak period of each separately active population firing within the cycle decreases towards lower (low beta or high alpha) bands. Thus, depending on the spatial distribution of the populations active in the working memory cycle, either gamma power may increase or else a “downshift” in measured frequencies (e.g., an increase in activity in or near the alpha band) may be expected with increasing working memory load. Neuronal networks may select for (coherent) oscillatory signals in the gamma range in particular [47], and increases in the spectral power in both gamma and the lower (e.g., alpha) frequency bands concomitant with increases in working memory load have been reported in a number of studies in humans [6, 23, 24].

Evidence of relationships between oscillations and the maintenance of working memory in humans has been obtained in numerous neurophysiological, imaging, and computational studies [6, 23–25, 27, 36, 48–54]. Oscillatory models have been developed both in the context of working memory [35, 55–57] and of binding [37, 58–60], and models – often with an eye towards image processing – have employed the distinction between bound and distinct objects as synchronous or asynchronous oscillations [37, 60–64]. These models tend to be spiking networks, appeal to cross-frequency coupling (e.g., theta-gamma codings), provide unrealistic connections (e.g., delayed self-inhibition for excitatory elements), use delays or constant inputs to produce persistent oscillatory activity, or employ structured architectures (e.g., using Hopfield networks, Hebbian rules, or pre-wired assemblies). In contrast, our model incorporates NMDA synapses to produce persistent activation and a simplified architecture (e.g., uniform all-to-all connectivity) that, importantly, allows any of the populations to be in- or out-of-phase with each other. Our model additionally has the advantage of being posed in a more mathematically tractable form than many other models, allowing us to analyze the dependence of states on different network parameters. Since we consider the ensemble activity through a mean-field model, we have also been able to explore more behaviors that the model provides, and have included several examples of interesting and relevant working memory and binding activities. In the present work, the networks are capable of producing the full range of oscillatory frequencies implicated in working memory and consistent with these other studies. However, we did not focus on particular frequency bands in terms of their implications for a working memory code, but rather on the role they can play in facilitating or providing a mechanism for the various characteristics of and essential functions underlying working memory.

We note that the mechanisms and facilitation of these working memory properties and functionality arise from, or map onto, the same general oscillatory dynamics generated in the network. That is, these dynamics provide the basic properties by which executive processes can control ongoing cognitive operations through the control of the contents of declarative or procedural working memory. The dynamics meet the potentially opposing demands of this control by providing a mechanism for quickly establishing and maintaining new structures or representations by both forming strong bindings that are resistant to interference and retaining the ability to rapidly dissolve those bindings in order to update the contents of working memory, or remove content from working memory that is no longer relevant. Thus, a primary result in this work is that binding via synchronization between populations that represent working memory elements, the number of arrangements of which appears limited only by the number of permutations of active populations, may be rapidly and stably formed as well as dissolved to form new structures from other populations that represent elements in long-term memory. However, while all arrangements of the active populations appear possible, different numbers of operations (e.g., activations or deactivations) are necessary to reach particular states from each other. This could be a factor involved in different lag times in the performance of different types of working memory tasks, as observed in numerous studies [17, 65–68].

We have not explicitly addressed timing issues or how the absolute order of memoranda are organized (for example, that two different, simultaneously-presented memoranda are recognized as separate, as opposed to being features of a single memorandum, or how an OP memorandum is recognized as the first in a sequence of repeating OP memoranda).

Rather, we have shown that the dynamics and combinatorial possibilities that are possible within working memory enable the different types of binding and other functionality and characteristics observed. Presumably, other circuitry (either feed-forward or recurrent) and mechanisms exist for these purposes. For example, the present work should be compatible with previous work addressing how the order of memoranda could be maintained through superimposing some external oscillatory signal in phase with the memory cycle (e.g., [27]). With respect to binding, many of the organization issues could be managed by either temporal (e.g., similar phase) or spatial (e.g., some feature map) proximity, or in the case of variable binding via some additional reference engine network as in Shastri et al. [42]. We also note that, while we have explored these networks within the context of the persistent activity observed in working memory dynamics, the presented behaviors would persist if we did not include NMDA dynamics, instead providing sustained drive to the excitatory and inhibitory populations (e.g., we show the same qualitative dynamics for such a reduced system with one population in Appendix A). Object differentiation and feature binding, as required in image segmentation, for example, may thus be modeled in our network using the same mechanisms described in the present results without consideration for persistent activity, so that simply removing the stimulus immediately quenches the activity of the selected populations, as in other work dealing with image analysis [61–64].

The ability of the networks in our model to bind distinct populations, unbind these populations and establish new bindings, and insert different numbers of new bound populations within a given cycle in working memory is made possible by the oscillatory dynamics. The combinatorial richness of binding possibilities that emerges provides a mechanistic description of working memory function that is consistent with other models incorporating, for example, feature binding or variable binding, and we examined examples of both of these in the present work. For the case of variable binding we have shown how the model can map onto and provide a mechanism consistent with inference networks that describe features of language or abstract reasoning. Here we showed how in a simple case these working memory networks can dynamically update and load new variable values into network nodes that bind together dynamically to carry out simple predicate calculi. We do not specify in the present case how specific nodes or variables would know to associate or bind, but rather that the dynamics exist for them to spontaneously do so. Presumably these associations would occur as a function of some type of map in which particular values would become associated on the basis of their closeness or specific connectivities established in long-term memory. Further work needs to be done with larger populations to develop these applications further. In the present work we do show that such associations emerge naturally, and could potentially provide the basis for a range of cognitive functions and phenomena, such as grammars in language (as touched on in the present results), or pattern completion and recognition. In this latter example, different memoranda that are dissimilar in various features or to some particular degree are recognized to have similarities that enable them to be grouped into a single type (i.e completing the pattern) by the presentation of the new input information. Such a mechanism could also be utilized in so-called chunking, in which disparate items are associated into some pattern, and reformed as a new single memorandum or representation, thereby dramatically decreasing working memory load.

Another of the most common binding situations is general coordination, which is already thought to involve synchronization of neuronal activity, and may also be accommodated within the present model. The availability of several different phase “slots”, associated here with working memory capacity, along with the ability of multiple populations to synchronize within each “slot”, allows for different phase-timing differences among different populations. Synchronous and asynchronous finger-tapping are two of the simplest examples of coordination, and are easily accommodated within the model (S and OP activity, respectively).

Most of the above binding situations were obtained with a uniform connectivity with deterministic dynamics. Breaking the network’s symmetry by allowing for heterogenous architectures or dynamics that are noisy or aperiodic may entail imperfect synchronizations that may be more susceptible to perturbations. In fact, such a mechanism may account for certain types of illusions, as errors involving binding activations and changes may occur with greater frequency. However, within reasonable bounds, qualitatively similar results may be obtained. For example, random sparse networks have also been shown to be able to synchronize [69], and we found that allowing for randomness in the all-to-all network did not qualitatively change our results. Indeed, heterogenous coupling may allow for additional relevant dynamics. As an example, with a distance-dependent coupling in the model with multiple populations, a stimulated population could interact with an already-active bound set of populations, deactivating one or more of the populations closest to it in feature space while leaving active and binding with the other, more distant, populations. This provides a possible mechanism for temporally changing the features of a memorandum, as an input can activate a quiescent population that may then both unbind features and itself bind with the continually-active populations. Such changes may also involve a rapid sequence of binding updates. Oscillatory dynamics facilitate such transitions by being able to switch off already-active populations, especially during the down portion of the oscillation, and activate the new populations. More complicated frequency couplings may be of interest and are also possible; e.g., in Fig 8B, we see that 2:1 locking ratios may be achieved. Such period-doublings that occur with distance-dependent coupling can also allow for, say, populations A and B to bind with population C by synchronizing in a pairwise, alternating fashion, leaving populations A and B unbound. Allowing for chaotic oscillatory dynamics (as seen to exist in E-I Wilson-Cowan networks in [70]) may allow for even richer synchronous relationships as in the spiking networks in Raffone and van Leeuwen [60].

## Appendix A: The *u, v, n* system

Here we examine the mechanisms by which persistent steady state and oscillatory behaviors arise in our model when the excitatory neurons are excited by a brief input stimulus. The NMDA dynamics ensure NMDA stays bounded within [0, 1] (given initial conditions in the same interval). The null surface for NMDA is sigmoidal 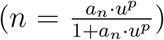, suggesting the possibility of bistability between a low state and a high state. Indeed, the NMDA allows for latch-like behavior of the system [28], so that low NMDA activity coincides with low *u* and *v*values, and high NMDA coincides with higher *u* and *v* values. Although Eq (1) describes a three-dimensional system, we may gain some intuition for the observed dynamics by fixing *n* at a constant level (since it evolves slowly and changes little compared to *u* and *v*) and examining the reduced *u* – *v* system. Thus, changing *n* in this planar system is equivalent to changing *θ*_e,i_ in tandem in the original system.

We therefore consider a projection of the autonomous version of the original system, Eq (1), where we take *n* as a parameter and forego the stimulus:

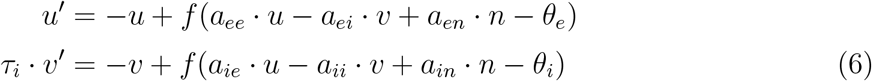

We may now examine the nullclines in Eq (6). As we see in Fig 11, for low *n* (Fig 11A) the *v*-nullcline intersects the left branch of the *u* nullcline, while for high *n* (Fig 11B), the *v*-nullcline intersects the middle branch of the *u*-nullcline. Thus, the low *n* case results in a stable fixed point, and the high *n* case results in a fixed point at higher *u* and *v* that may be stable or unstable, depending on τ_i_. For low τ_i_ (e.g., *τ*_i_ = 1), this high fixed point is stable; for higher τ_i_, a stable oscillation emerges, as suggested in the example trajectories in Fig 11B. Therefore, varying *n* (still as a parameter) allows such a two-dimensional system to switch between low and high activity states that can be either steady-state or oscillatory.

**Fig 11.**
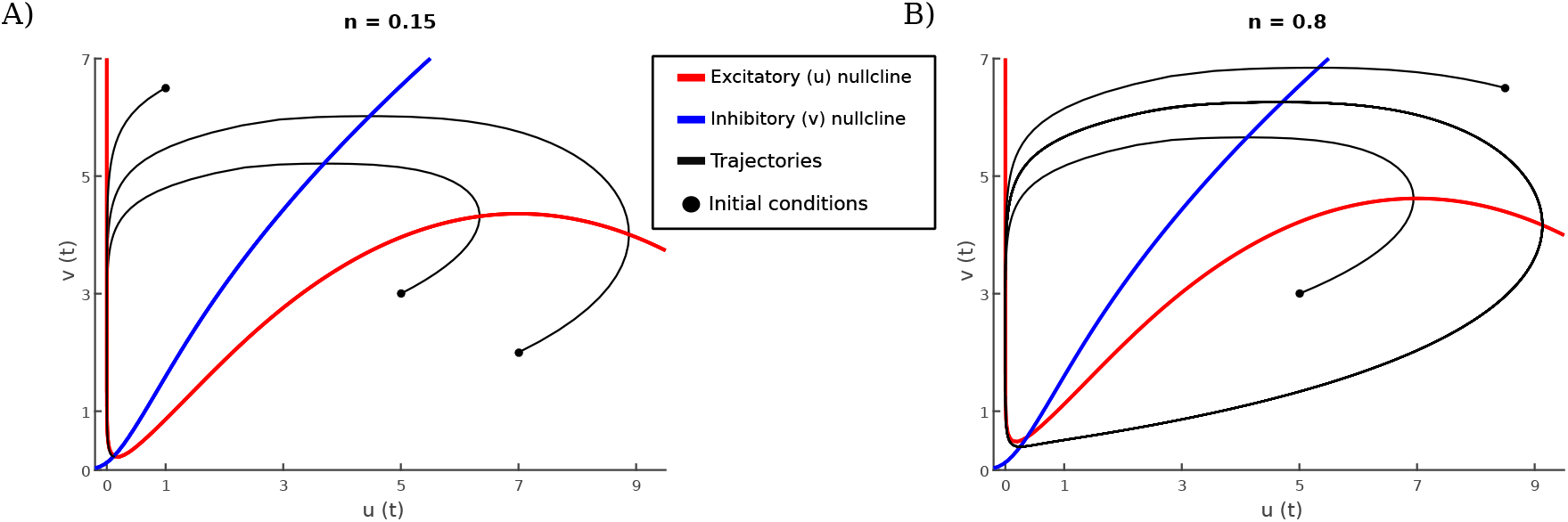
Nullclines and example trajectories for Eq (6). Parameter values are as given in section *2*. (A) For lower NMDA (*n*) values, the v-nullcline intercepts the left branch of the u-nullcline, resulting in a stable fixed point. (B) For higher NMDA values, the v-nullcline intercepts the middle branch of the u-nullcline; the fixed point is stable for low *τ*_*i*_ values and unstable for larger *τ*_*i*_ values. Here, *τ*_*i*_ = 12, so that the fixed point is unstable and the system has a stable limit cycle, as suggested by the example trajectories.

In the full three-dimensional system, we set up our *u* and *v* nullsurfaces so that their low-*n* and high-*n* cross-sections intersect exactly as the corresponding 2-dimensional nullclines do. As the NMDA nullsurface is sigmoidal, it can act as a dynamic latch, so that once *n* is excited enough (via the *s* (*t*) → *u* → *n* path), the system is attracted to the high state. The existence of bistability is, of course, dependent on system parameters; for example, we see that two stable fixed points exist for a range of *a*_*ei*_ values in Fig 12A. As *τ*_i_ increases, the high fixed point destabilizes and a limit cycle is born via a subcritical Hopf bifurcation (Fig 12B).

**Fig 12.**
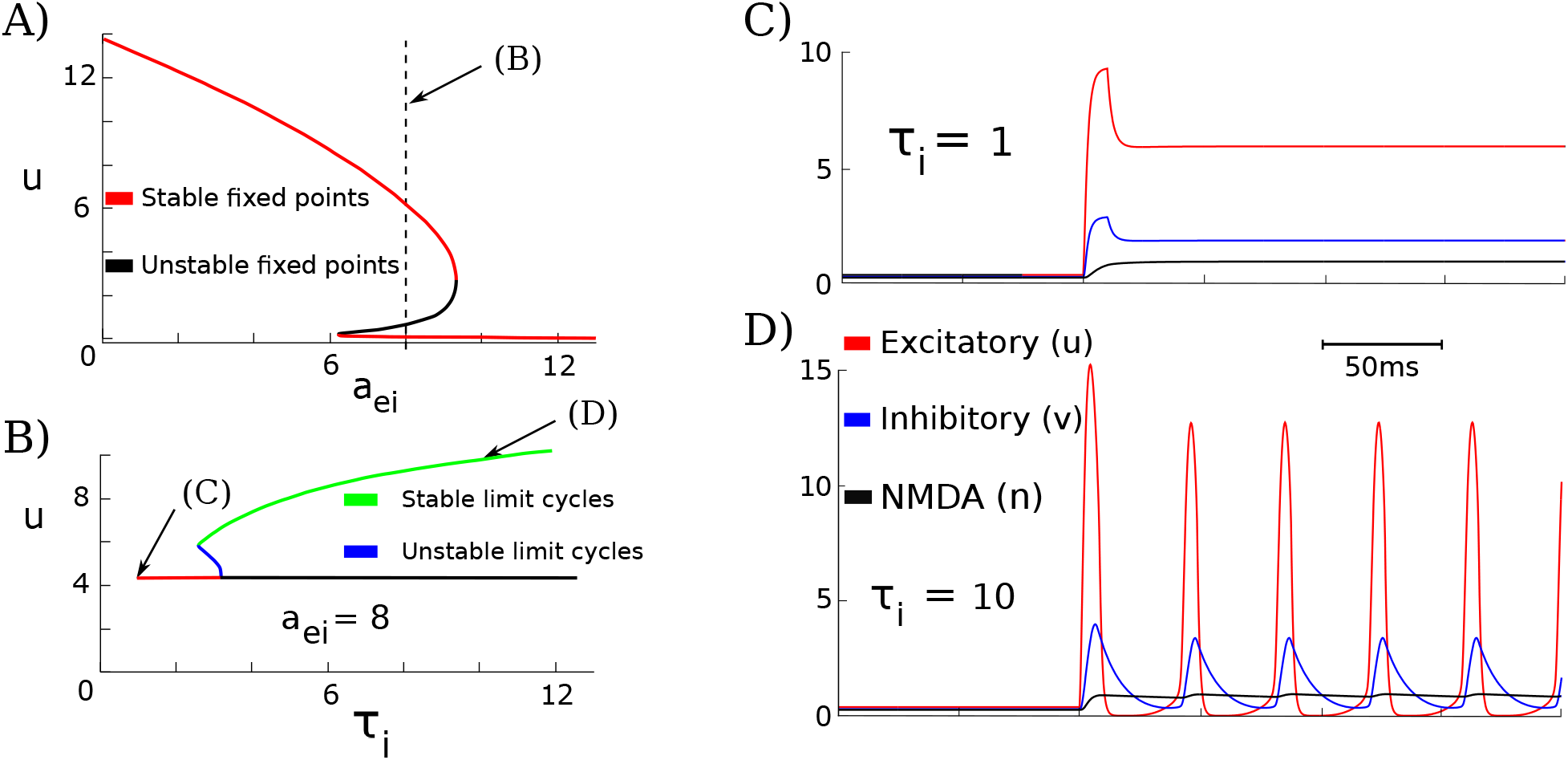
Bifurcations and dynamics for one population for Eq (1). (A) There are open sets of parameters, such as *a_ei_*, that allow for bistability between low states and high states. Here, *τ*_*i*_ = 1. (B) Increasing *τ_i_* leads to a s ubcritical Hopf bifurcation of the high fixed point. (C), (D) display the dynamics for *a*_*ei*_ = 8 and different *τ*_*i*_, leading to bistability of fixed points (C) or limit cycles (D).

We may gain further intuition for how the separation of timescales allows for oscillations. Once the excitatory stimulus kicks the AMPA population sufficiently, it begins an excursion around phase space. In turn, it excites the inhibitory population, which then chases the AMPA population, curtailing its growth. As the inhibitory timescale is slower than that of the excitatory population (*τ*_*i*_ < *τ*_*e*_ > 1), the inhibition eventually wins out, quenching the activity of the AMPA population, as we see in Fig 12D. This is why the NMDA population is important: The NMDA is excited by the AMPA, ut decays much slower than the inhibition (*τ*_*n*_ < *τ*_*i*_), allowing the NMDA to outlast the upstroke of the inhibition. By doing so, the NMDA can then re-excite the AMPA population once the inhibition is sufficiently low, producing the observed wavetrain in Fig 12D.

Thus, if *τ_n_* is too small, the NMDA population will increase rapidly with AMPA, but will then decay too quickly to maintain the activity of the AMPA population after the inhibitory population has quenched it. Clearly the same problem will occur if *τ*_*i*_ is too large. Therefore, the ratio 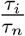 must be small enough. Finally, we note that for *τ*_*i*_ very small, the high fixed point is stable and thus acts as an attractor. This does not allow for the initial phase space excursion that occurs in the oscillatory case to take place at all, as seen in Fig 12C. Hence, we now have some intuition to understand the general shapes of the bifurcation curves in Fig 5B, 5C, and 5D.

## Appendix B: Spiking model comparison

Consider the mean first passage time for a noise-driven quadratic integrate-and-fire (QIF) model,

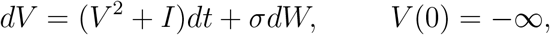

where σ is the noise and *I* is the drive. The expected time *T*(*I*, σ) for *V*(*t*) to reach infinity leads to the expected firing rate, *v*(I, σ) = 1/T. For the zero noise case, 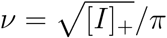, where [*I*]_+_ is the positive part of *I*. With noise, this rate can be closely approximated by the nonlinearity:

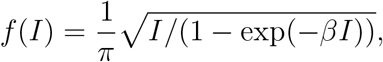

where *β* is chosen to best fit for a given σ [32].

With this in mind, we created a population of noisy QIF neurons in order to justify our simple three-variable model. The network has the form:

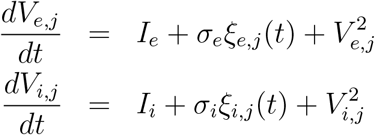

where *ξ* are independent white noise processes and the drives are:

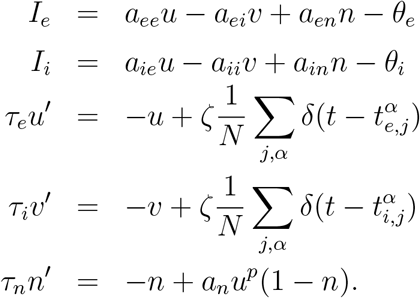

Each time *V* crosses +∞ (spike), it is reset to –*∞* and the appropriate variable, *u* or *v*, is incremented (as represented by the sums of delta functions). We set *N* = 200 for both the E and I populations and solved these equations with Euler’s method and a time step of 0.005. We set the the spike value to +100 and the reset to −100. In order to match the firing rate curve of a single population, we set the noise amplitude to *σ*_*e,i*_ = 3, which matched *β* = 1 reasonably well. The scale factor *ζ* is included since we do not divide our firing rate function by *π*. Thus, the scale factor should be set to be *π*; we found that if we lowered it a little bit to *ζ* = 2.9, we got better matching. Fig 13 shows the result of a simulation where the parameters for the spiking model are exactly the same as in the simple (*u*, *v*, *n*) system in Eq (1). Here we have started *n* at a high value to obtain the oscillatory state. This demonstrates that we can create a spiking network that has the same properties as our simple mean field network.

**Fig 13.**
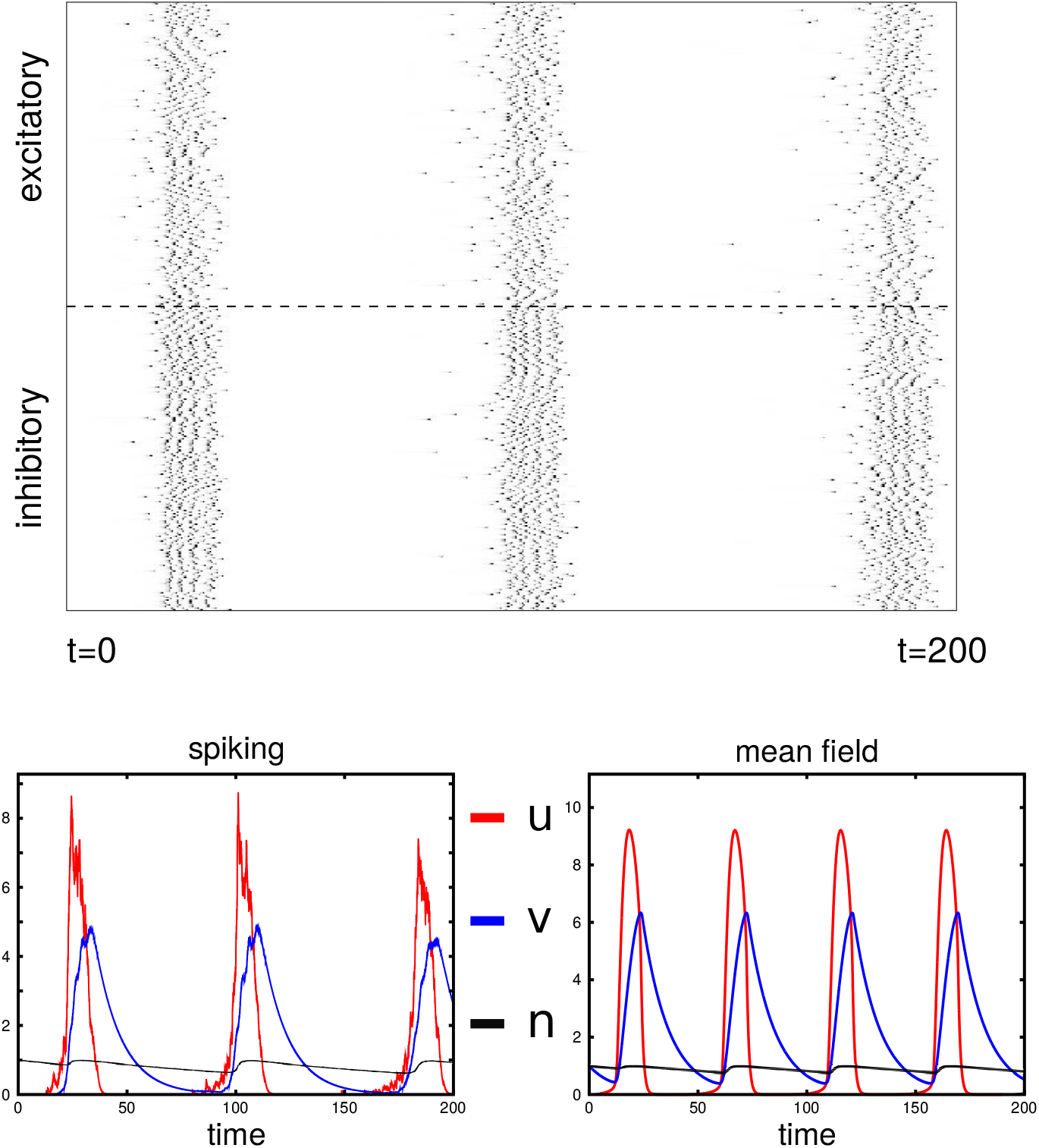
Spiking simulations and comparison to the mean field equations. Top: Raster plot showing the spikes of the 200 excitatory and 200 inhibitor y neurons over a little more than one cycle. Bottom: The triplet *(u,v, n)* for the spiking network (on the left) compared to the mean field behavior (right).

## Appendix C: Weak coupling analysis

Using the standard parameters for a single local *u* – *v* – *n* circuit, we can perform a weak coupling analysis (WCA) around the limit cycle solution that appears in the up state. That is, since the coefficients that we use tend to be small compared to the self coupling, the WCA can provide some insights into the possible patterns when *m* groups are oscillating. The results should hold for any size network, since the “down” circuits contribute very little to the interactions in the groups that are oscillating. To perform the WCA, we first see that

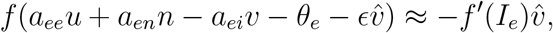

where *I*_*e*_ = *a*_*ee*_*u* + *a*_*en*_*n* – *a*_*ei*_*v* – *θ*_*e*_, and with similar terms for other types of coupling. Recall that in WCA, we reduce

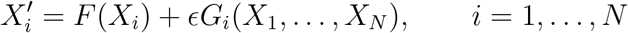

to a set of equations of the form:

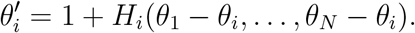

To achieve this reduction, we assume there is a *T*–periodic limit cycle solution, *U’* = *F*(*U*), and let *Z*(*t*) be the unique solution (adjoint solution) to *Z’* = –(*D*_x_*F*(*U*))^T^, with *U’*(*t*) · *Z*(*t*) = 1. The phase interaction functions *H*_*i*_ are defined as

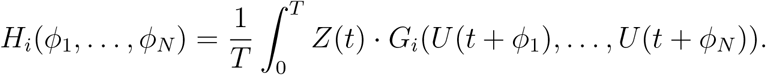

Since all coupling is summed up, we only need to compute the interaction with one other circuit. Thus, we need to compute the the adjoint solution and the basic limit cycle. Letting (*u*(*t*), *v*(*t*), *n*(*t*)) be the basic limit cycle for the isolated population and (*u*^*\**^ (*t*), *v*^*\**^(*t*), *n*^*\**^(*t*)) be the corresponding adjoint solution, we compute the following four interaction functions:

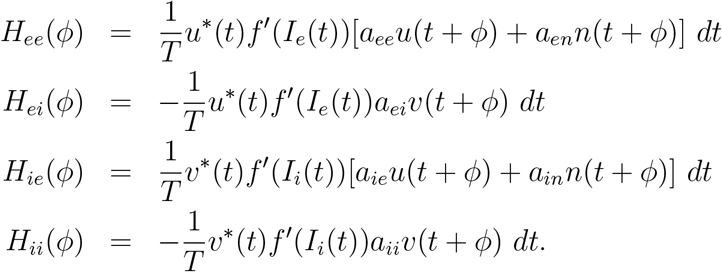

Once we have the interaction functions, we can study the dynamics of the oscillators when weakly coupled. We form the composite function:

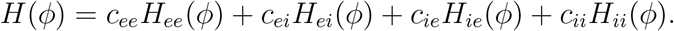

The coupled phase equations satisfy

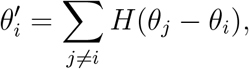

where *i* varies from 1 to *N* active groups. To determine the locked patterns, we reduce the dimension to *N* – 1 by setting *θ*_1_ = 0 and subtracting *θ*_1_ from the remaining *N* – 1 equations:

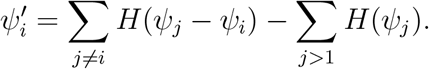

Here, *ψ*_*i*_ represents the phase relative to *θ*_1_. Stable fixed points of this (*N* –1) – dimensional system correspond to the attracting dynamics of the weakly coupled system. Table 1 summarizes the attractors for up to 4 active groups when each of the individual coupling terms are set to 1 and the rest are set to zero. We note that the behavior of EE coupling serves mainly to synchronize and that the EI and IE coupling have similar behavior. Small amounts of EE coupling in addition to EI coupling (what we use in the full model) behave like the EI coupling alone as long as the EE coupling is not too big. For example, with EI coupling and three active groups, WCA predicts that there will be synchrony (S), a splay state (L), and a clustered state (C) with two oscillators synchronized and the third out of phase (see Fig 14). Since the coupling is all-to-all and symmetric, all possible permutations of the attractors occur.

**Fig 14.**
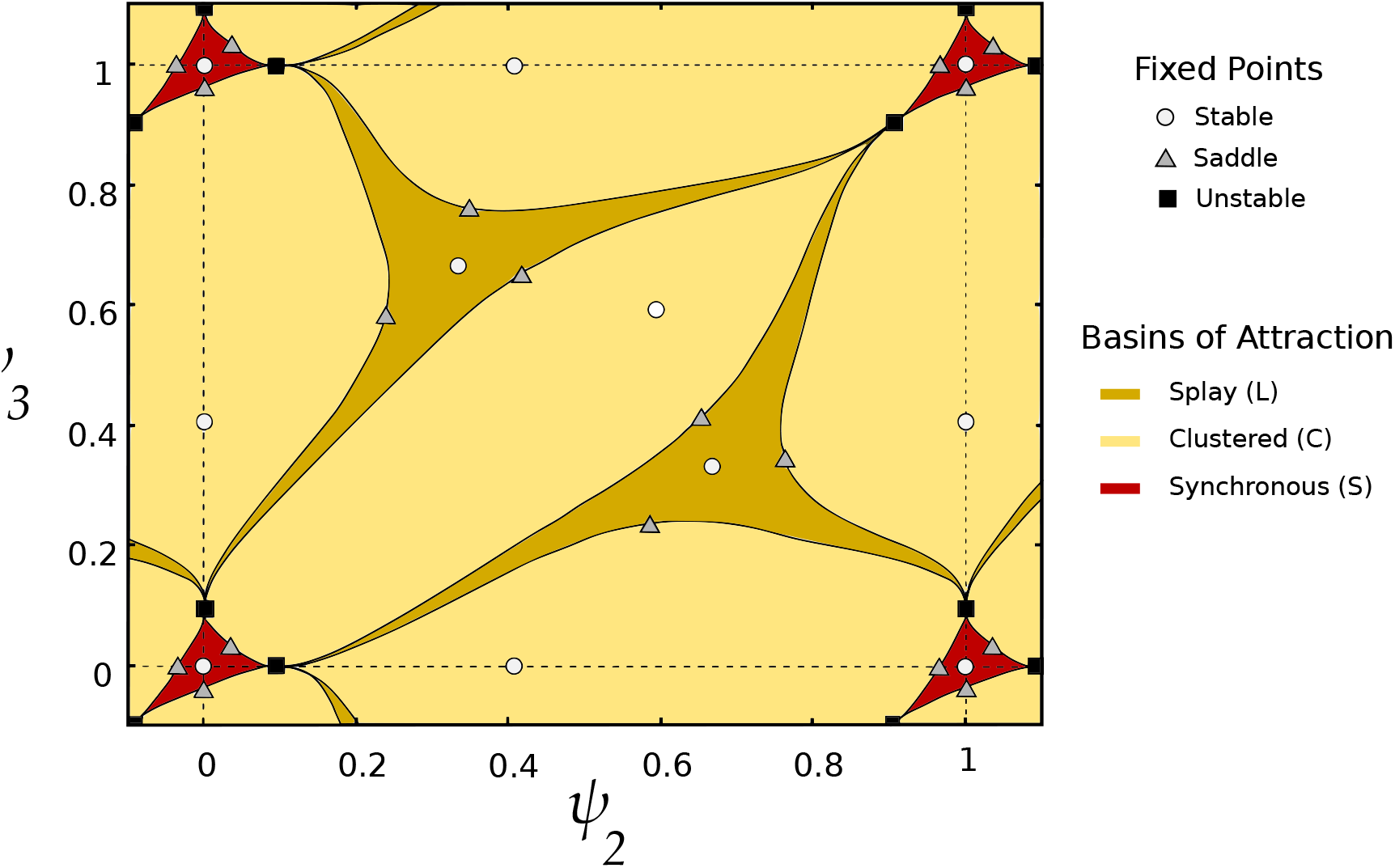
Basins of attraction for weak EI coupling with 3 active populations. When three populations are active (note, *ψ*_*i*_ is the phase of population *i* relative to *θ*_1_), splay (L), clustered (C), and synchronous (S) states all have open sets as basins of attraction, as we found in the full model. These basins are defined by the stable manifolds of saddle points, as shown.

This analysis explains many of the interactions we see with active populations. We remark that the analysis is only valid when the coupling is weak enough, but it still manages to include many of the attractors that are seen when there are two or more active populations. We finally note that for 5 or more active populations we only see various clustered states and synchrony with weak coupling; we do not see any splay states. Thus, at least when the coupling is weak, there is limited capacity in the network.

## References

1. Fuster JM, Alexander GE, et al. Neuron activity related to short-term memory. Science. 1971;173(3997):652–654.

2. Wang XJ. Synaptic basis of cortical persistent activity: the importance of NMDA receptors to working memory. Journal of Neuroscience. 1999;19(21):9587–9603.

3. Fuster JM, Bodner M, Kroger JK. Cross-modal and cross-temporal association in neurons of frontal cortex. Nature. 2000;405(6784):347.

4. Pesaran B, Pezaris JS, Sahani M, Mitra PP, Andersen RA. Temporal structure in neuronal activity during working memory in macaque parietal cortex. Nature Neuroscience. 2002;5(8):805–812.

5. Shafi M, Zhou Y, Quintana J, Chow C, Fuster J, Bodner M. Variability in neuronal activity in primate cortex during working memory tasks. Neuroscience. 2007;146(3):1082–1108.

6. Ku Y, Bodner M, Zhou YD. Prefrontal cortex and sensory cortices during working memory: quantity and quality. Neurosci Bull. 2015;31(2):175–182.

7. Cowan N. The magical number 4 in short-term memory: A reconsideration of mental storage capacity. Behavioral and Brain Sciences. 2001;24(1):87–114.

8. Baddeley A. Working memory: looking back and looking forward. Nature Reviews Neuroscience. 2003;4(10):829.

9. Luck SJ, Vogel EK. The capacity of visual working memory for features and conjunctions. Nature. 1997;390(6657):279.

10. Vogel EK, Woodman GF, Luck SJ. Storage of features, conjunctions, and objects in visual working memory. Journal of Experimental Psychology: Human Perception and Performance. 2001;27(1):92.

11. Todd JJ, Marois R. Capacity limit of visual short-term memory in human posterior parietal cortex. Nature. 2004;428(6984):751.

12. Feldman J. The neural binding problem(s). Cognitive neurodynamics. 2013;7(1):1–11.

13. Von Der Malsburg C. The correlation theory of brain function. In: Models of neural networks. Springer; 1994. p. 95–119.

14. Singer W, Gray CM. Visual feature integration and the temporal correlation hypothesis. Annual review of neuroscience. 1995;18(1):555–586.

15. Singer W. Neuronal synchrony: a versatile code for the definition of relations? Neuron. 1999;24(1):49–65.

16. Engel AK, König P, Singer W. Direct physiological evidence for scene segmentation by temporal coding. Proceedings of the National Academy of Sciences. 1991;88(20):9136–9140.

17. Kessler Y, Meiran N. All updateable objects in working memory are updated whenever any of them are modified: Evidence from the memory updating paradigm. Journal of Experimental Psychology: Learning, Memory, and Cognition. 2006;32(3):570.

18. Fries P, Reynolds JH, Rorie AE, Desimone R. Modulation of oscillatory neuronal synchronization by selective visual attention. Science. 2001;291(5508):1560–1563.

19. Cowan N, Blume CL, Saults JS. Attention to attributes and objects in working memory. Journal of Experimental Psychology: Learning, Memory, and Cognition. 2013;39(3):731.

20. Hardman KO, Cowan N. Remembering complex objects in visual working memory: Do capacity limits restrict objects or features? Journal of Experimental Psychology: Learning, Memory, and Cognition. 2015;41(2):325.

21. Klimesch W. EEG alpha and theta oscillations reflect cognitive and memory performance: a review and analysis. Brain research reviews. 1999;29(2-3):169–195.

22. Başar E, Başar-Eroglu C, Karakaş S, Schürmann M. Gamma, alpha, delta, and theta oscillations govern cognitive processes. International journal of psychophysiology. 2001;39(2-3):241–248.

23. Howard MW, Rizzuto DS, Caplan JB, Madsen JR, Lisman J, Aschenbrenner-Scheibe R, et al. Gamma oscillations correlate with working memory load in humans. Cerebral cortex. 2003;13(12):1369–1374.

24. Jensen O, Gelfand J, Kounios J, Lisman JE. Oscillations in the alpha band (9–12 Hz) increase with memory load during retention in a short-term memory task. Cerebral cortex. 2002;12(8):877–882.

25. Roux F, Uhlhaas PJ. Working memory and neural oscillations: alpha–gamma versus theta–gamma codes for distinct WM information? Trends in cognitive sciences. 2014;18(1):16–25.

26. Verduzco-Flores S, Ermentrout B, Bodner M. From working memory to epilepsy: Dynamics of facilitation and inhibition in a cortical network. Chaos: An Interdisciplinary Journal of Nonlinear Science. 2009;19(1):015115.

27. Lisman J. Working memory: the importance of theta and gamma oscillations. Current Biology. 2010;20(11):R490–R492.

28. Lisman JE, Fellous JM, Wang XJ. A role for NMDA-receptor channels in working memory. Nature neuroscience. 1998;1(4):273–275.

29. Adler CM, Goldberg TE, Malhotra AK, Pickar D, Breier A. Effects of ketamine on thought disorder, working memory, and semantic memory in healthy volunteers. Biological psychiatry. 1998;43(11):811–816.

30. Krystal JH, Abi-Saab W, Perry E, D’Souza DC, Liu N, Gueorguieva R, et al. Preliminary evidence of attenuation of the disruptive effects of the NMDA glutamate receptor antagonist, ketamine, on working memory by pretreatment with the group II metabotropic glutamate receptor agonist, LY354740, in healthy human subjects. Psychopharmacology. 2005;179(1):303–309.

31. Compte A, Brunel N, Goldman-Rakic PS, Wang XJ. Synaptic mechanisms and network dynamics underlying spatial working memory in a cortical network model. Cerebral Cortex. 2000;10(9):910–923.

32. Ermentrout GB, Terman DH. Mathematical foundations of neuroscience. vol. 35. Springer Science & Business Media; 2010.

33. Oberauer K. Interference between storage and processing in working memory: Feature overwriting, not similarity-based competition. Memory & Cognition. 2009;37(3):346–357.

34. Bancroft TD, Servos P, Hockley WE. Mechanisms of interference in vibrotactile working memory. PLoS One. 2011;6(7):e22518.

35. Lisman JE, Idiart MA. Storage of 7+/-2 short-term memories in oscillatory subcycles. Science. 1995;267(5203):1512–1515.

36. Lisman JE, Jensen O. The theta-gamma neural code. Neuron. 2013;77(6):1002–1016.

37. Raffone A, Wolters G. A cortical mechanism for binding in visual working memory. Journal of Cognitive Neuroscience. 2001;13(6):766–785.

38. Worden MS, Foxe JJ, Wang N, Simpson GV. Anticipatory biasing of visuospatial attention indexed by retinotopically specific-band electroencephalography increases over occipital cortex. J Neurosci. 2000;20(RC63):1–6.

39. Klimesch W, Sauseng P, Hanslmayr S. EEG alpha oscillations: the inhibition–timing hypothesis. Brain research reviews. 2007;53(1):63–88.

40. Klimesch W. Alpha-band oscillations, attention, and controlled access to stored information. Trends in cognitive sciences. 2012;16(12):606–617.

41. Jensen O, Bonnefond M, VanRullen R. An oscillatory mechanism for prioritizing salient unattended stimuli. Trends in cognitive sciences. 2012;16(4):200–206.

42. Shastri L, Ajjanagadde V. From simple associations to systematic reasoning: A connectionist representation of rules, variables and dynamic bindings using temporal synchrony. Behavioral and brain sciences. 1993;16(3):417–451.

43. Wendelken C, Shastri L. Multiple instantiation and rule mediation in SHRUTI. Connection Science. 2004;16(3):211–217.

44. Sakai K, Rowe JB, Passingham RE. Active maintenance in prefrontal area 46 creates distractor-resistant memory. Nature neuroscience. 2002;5(5):479.

45. Buzsáki G, Wang XJ. Mechanisms of gamma oscillations. Annual review of neuroscience. 2012;35:203–225.

46. Whittington MA, Traub R, Kopell N, Ermentrout B, Buhl E. Inhibition-based rhythms: experimental and mathematical observations on network dynamics. International journal of psychophysiology. 2000;38(3):315–336.

47. Bürgers C, Kopell NJ. Gamma oscillations and stimulus selection. Neural computation. 2008;20(2):383–414.

48. Voytek B, Canolty RT, Shestyuk A, Crone NE, Parvizi J, Knight RT. Shifts in gamma phase–amplitude coupling frequency from theta to alpha over posterior cortex during visual tasks. Frontiers in human neuroscience. 2010;4.

49. Tallon-Baudry C, Bertrand O, Peronnet F, Pernier J. Induced *γ*-band activity during the delay of a visual short-term memory task in humans. Journal of Neuroscience. 1998;18(11):4244–4254.

50. Tallon C, Bertrand O, Bouchet P, Pernier J. Gamma-range activity evoked by coherent visual stimuli in humans. European Journal of Neuroscience. 1995;7(6):1285–1291.

51. Roux F, Wibral M, Mohr HM, Singer W, Uhlhaas PJ. Gamma-band activity in human prefrontal cortex codes for the number of relevant items maintained in working memory. Journal of Neuroscience. 2012;32(36):12411–12420.

52. Medendorp WP, Kramer GF, Jensen O, Oostenveld R, Schoffelen JM, Fries P. Oscillatory activity in human parietal and occipital cortex shows hemispheric lateralization and memory effects in a delayed double-step saccade task. Cerebral cortex. 2006;17(10):2364–2374.

53. Kaiser J, Rahm B, Lutzenberger W. Temporal dynamics of stimulus-specific gamma-band activity components during auditory short-term memory. Neuroimage. 2009;44(1):257–264.

54. Lutzenberger W, Ripper B, Busse L, Birbaumer N, Kaiser J. Dynamics of gamma band activity during an audiospatial working memory task in humans. Journal of neuroscience. 2002;22(13):5630–5638.

55. Wang D, Buhmann J, von der Malsburg C. Pattern segmentation in associative memory. Neural Computation. 1990;2(1):94–106.

56. Horn D, Usher M. Parallel activation of memories in an oscillatory neural network. Neural computation. 1991;3(1):31–43.

57. Winder RK, Reggia JA, Weems SA, Bunting MF. An oscillatory Hebbian network model of short-term memory. Neural Computation. 2009;21(3):741–761.

58. König P, Schillen TB. Stimulus-dependent assembly formation of oscillatory responses: I. Synchronization. Neural Computation. 1991;3(2):155–166.

59. Sompolinsky H, Golomb D, Kleinfeld D. Global processing of visual stimuli in a neural network of coupled oscillators. Proceedings of the National Academy of Sciences. 1990;87(18):7200–7204.

60. Raffone A, van Leeuwen C. Dynamic synchronization and chaos in an associative neural network with multiple active memories. Chaos: An Interdisciplinary Journal of Nonlinear Science. 2003;13(3):1090–1104.

61. Terman D, Wang D. Global competition and local cooperation in a network of neural oscillators. Physica D: Nonlinear Phenomena. 1995;81(1-2):148–176.

62. Meier M, Haschke R, Ritter HJ. Perceptual grouping through competition in coupled oscillator networks. Neurocomputing. 2014;141:76–83.

63. Yu G, Slotine JJ. Visual grouping by neural oscillator networks. IEEE Transactions on neural Networks. 2009;20(12):1871–1884.

64. Breve FA, Zhao L, Quiles MG, Macau EE. Chaotic phase synchronization and desynchronization in an oscillator network for object selection. Neural Networks. 2009;22(5-6):728–737.

65. Oberauer K. Declarative and procedural working memory: common principles, common capacity limits? Psychologica Belgica. 2010;50(3-4):3–4.

66. Garavan H. Serial attention within working memory. Memory & cognition. 1998;26(2):263–276.

67. Oberauer K. Selective attention to elements in working memory. Experimental psychology. 2003;50(4):257.

68. Kessler Y, Meiran N. Two dissociable updating processes in working memory. Journal of Experimental Psychology: Learning, Memory, and Cognition. 2008;34(6):1339.

69. Börgers C, Kopell N. Synchronization in networks of excitatory and inhibitory neurons with sparse, random connectivity. Neural computation. 2003;15(3):509–538.

70. Borisyuk GN, Borisyuk RM, Khibnik AI, Roose D. Dynamics and bifurcations of two coupled neural oscillators with different connection types. Bulletin of mathematical biology. 1995;57(6):809–840.

